# Impaired axonal transport at the optic nerve head contributes to neurodegeneration in a novel Cre-inducible mouse model of myocilin glaucoma

**DOI:** 10.1101/2024.09.18.613712

**Authors:** Balasankara Reddy Kaipa, Ramesh Kasetti, Yogapriya Sundaresan, Linya Li, Sam Yacoub, J. Cameron Millar, William Cho, Dorota Skowronska-Krawczyk, Prabhavathi Maddineni, Krzysztof Palczewski, Gulab S. Zode

**Author notes:** Corresponding author: Gulab S. Zode, PhD 829 Health Sciences Rd, Irvine, CA 92617.

## Abstract

Elevation of intraocular pressure (IOP) due to trabecular meshwork (TM) dysfunction, leading to neurodegeneration, is the pathological hallmark of primary open-angle glaucoma (POAG). Impaired axonal transport is an early and critical feature of glaucomatous neurodegeneration. However, a robust mouse model that replicates these human POAG features accurately has been lacking. We report the development and characterization of a novel Cre-inducible mouse model expressing a DsRed-tagged Y437H mutant of human myocilin (*Tg.CreMYOC^Y^*^437^*^H^*). A single intravitreal injection of HAd5-Cre induced selective MYOC expression in the TM, causing TM dysfunction, reducing outflow facility, and progressively elevating IOP in *Tg.CreMYOC^Y^*^437^*^H^* mice. Sustained IOP elevation resulted in significant retinal ganglion cell (RGC) loss and progressive axonal degeneration in Cre-induced *Tg.CreMYOC^Y^*^437^*^H^* mice. Notably, impaired anterograde axonal transport was observed at the optic nerve head before RGC degeneration, independent of age, indicating that impaired axonal transport contributes to RGC degeneration in *Tg.CreMYOC^Y^*^437^*^H^* mice. In contrast, axonal transport remained intact in ocular hypertensive mice injected with microbeads, despite significant RGC loss. Our findings indicate that Cre-inducible *Tg.CreMYOC^Y^*^437^*^H^*mice replicate all glaucoma phenotypes, providing an ideal model for studying early events of TM dysfunction and neuronal loss in POAG.

## Introduction

Glaucoma is a group of multifactorial neurodegenerative diseases characterized by progressive optic neuropathy. It is the second leading cause of irreversible vision loss, affecting more than 70 million people affected worldwide ^1, 2^, and this number is estimated to increase to 112 million by the year 2040 ^3^. Primary open angle glaucoma (POAG) is the most common form of glaucoma, accounting for approximately 70% of all the cases ^1^. POAG is characterized by progressive loss of retinal ganglion cell (RGC) axons and irreversible loss of vision^1, 2^. Impaired axonal transport at the optic nerve head (ONH) is implicated as an early pathological event associated with glaucomatous neurodegeneration^4–6^. Elevated intraocular pressure (IOP) is a major risk factor, and the only treatable risk factor for POAG ^7^. The trabecular meshwork (TM), a molecular sieve-like structure, regulates IOP by constantly adjusting the resistance to aqueous humor (AH) outflow. In POAG, increased resistance to AH outflow elevates IOP leading to neurodegeneration^8–10^. This increase in outflow resistance is associated with TM dysfunction^11–15^. Most of the current treatment strategies for POAG do not target the underlying pathology of the TM and RGCs, and vision loss continues to progress in some patients^16^. To understand the pathological mechanisms of TM dysfunction/IOP elevation and glaucomatous neurodegeneration, there is an unmet need to develop a simple and reliable animal model that can faithfully replicate all features of POAG.

Currently, several mouse models of ocular hypertension (OHT) are utilized to study the pathophysiology of glaucomatous neurodegeneration^17, 18^. These include DBA/2J mice and inducible mouse models that physically block AH outflow through the TM leading to OHT and neuronal loss^19–24^. While these models have provided important mechanistic insights, they do not allow the study of TM pathology, as they elevate IOP by physically blocking the outflow pathway ^21–23^. Thus, these models do not truly represent the human-POAG phenotype where the AH outflow pathway is open. Importantly, these models destroy the TM, which can induce acute and sudden IOP-induced glaucomatous neurodegeneration. Moreover, these models are often technically challenging to replicate in the laboratory settings, and they exhibit variable phenotypes and other confounding features such as ocular inflammation. Developing a mouse model mimicking a known genetic cause of human POAG represents an ideal strategy to understand the pathophysiology of POAG.

Mutations in myocilin (*MYOC*) are the most common genetic cause of POAG and are a significant contributor to juvenile-onset OAG (JOAG) ^9, 25, 26^ ^27^. MYOC-associated JOAG is a more aggressive form of glaucoma, characterized by high IOP in young children and rapid progression to vision loss^9, 26^. Mouse models with these genetic alterations in the *MYOC* gene mimicking human POAG have proved to be invaluable tools for understanding the pathogenesis of POAG and designing treatment strategies^28–30^. Previously, we have developed a transgenic mouse model (*Tg-MYOC^Y^*^437^*^H^*) of myocilin POAG by random genomic insertion of the human mutant myocilin and demonstrated that *Tg-MYOC^Y^*^437^*^H^* mice develop glaucoma phenotypes closely resembling those seen in POAG patients with the Y437H *MYOC* mutation^30^. However, *Tg-MYOC^Y^*^437^*^H^* mice presented a few drawbacks including 1) possible unknown gene mutations and multiple copies of the transgene due to random integration; 2) a mild phenotype on pure a C57BL/6J background limiting the study of glaucomatous neurodegeneration; 3) possible silencing of the transgene upon subsequent breeding; and 4) lack of specific antibodies to detect myocilin in the mouse TM. To overcome these limitations, we utilized a TARGATT site-specific knock in strategy^31^ to generate transgenic mice expressing human mutant *MYOC*. This technology employs serine integrase, PhiC31 (ΦC31) to insert a single copy of gene of interest into a preselected intergenic and transcriptionally active genomic locus (H11), which has been engineered with a docking site. This allows stable and single site-specific transgene integration. Since mutant MYOC is toxic to TM cells^30, 32^, we exploited the Cre-lox system to develop an inducible mouse model in which mutant MYOC is expressed in the tissue of interest only upon Cre expression. Here, we report the development and characterization of a Cre-inducible transgenic mouse line expressing the DsRed-tagged Y437H mutant of human myocilin (*Tg.CreMYOC^Y^*^437^*^H^*). We further utilized this model to investigate early events of glaucomatous TM dysfunction and neurodegeneration. In contrast to the microbead-occlusion model of OHT, in which axonal transport remains intact despite significant RGC loss, we observed that sustained IOP elevation in the *Tg.CreMYOC^Y^*^437^*^H^* mice significantly impairs axonal transport at the ONH prior to RGC soma and axonal loss.

## Results

### Generation of an inducible mouse model of myocilin POAG

Using TARGATT site-specific knock-in strategy, we developed a Cre-inducible transgenic mice that express the DsRed-tagged Y437H mutant of human *MYOC* (referred as *Tg.CreMYOC^Y^*^437^*^H^*). Under normal conditions, the mice do not express the human mutant-*MYOC* gene. Expression of Cre recombinase, however, leads to removal of a Stop cassette and consequent expression of the DsRed-fused mutant *MYOC* (Figure 1A). A single copy of transgene is inserted into a preselected intergenic and transcriptionally active genomic locus (H11) that has been engineered with a docking site for stable and site-specific transgene integration. To confirm a site-specific knock-in of transgene at the H11 site, we first performed PCR using primers specific to the integration site (Figure SI-1A, Supplemental Information), which demonstrated a stable integration of transgene in one founder line. These founder mice were further bred with C57BL/6J mice, and offspring were utilized for subsequent studies. For routine genotyping, primers specific to DsRed were utilized, which confirmed the presence of the transgene (Figure SI-1B).

**Figure 1.**
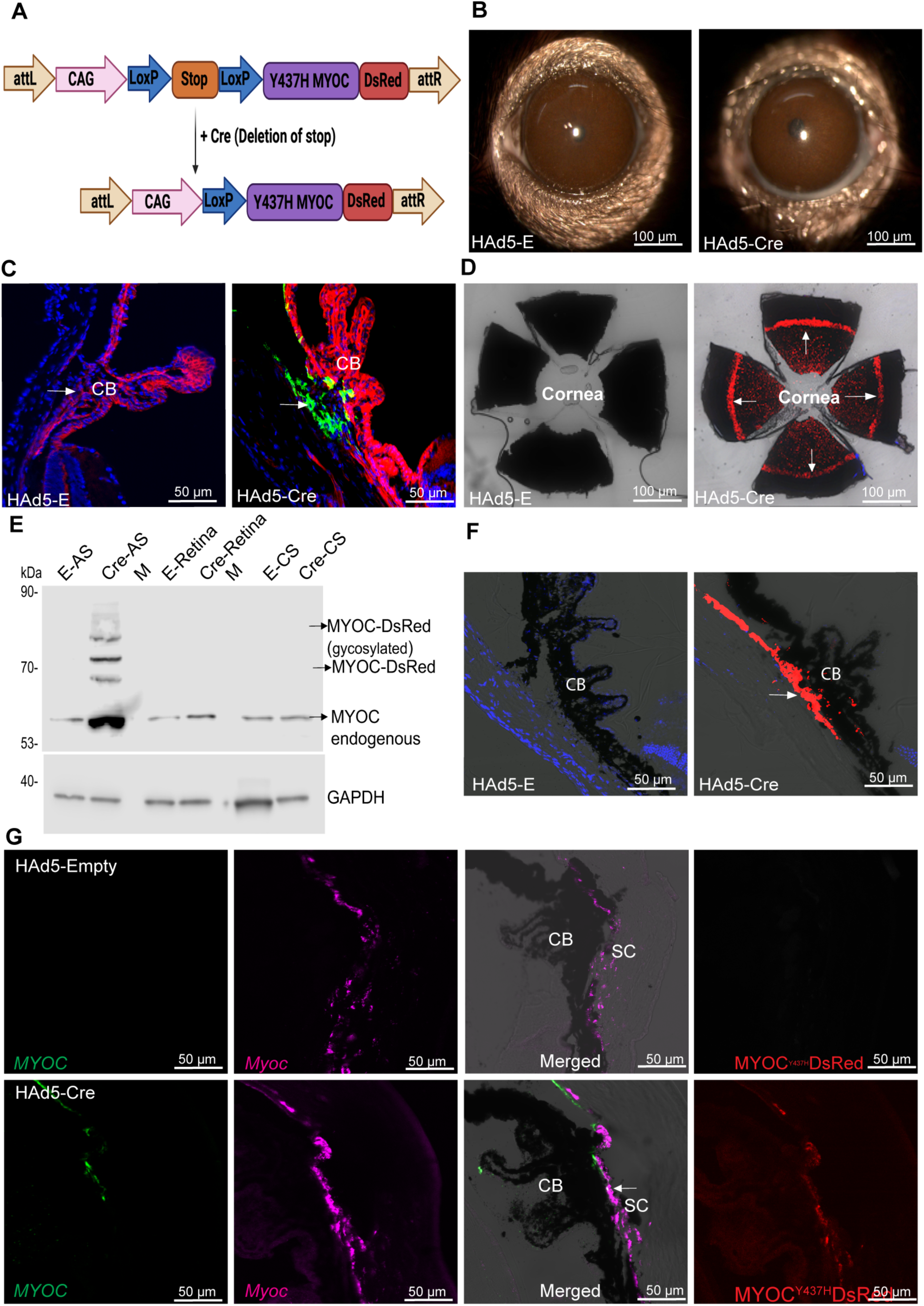
HAd5-Cre recombinase induces mutant MYOC in mouse TM. (**A**) Design Strategy: *Tg.CreMYOC^Y^*^437^*^H^* mice were engineered using a TARGATT gene knock in strategy in which DsRed-tagged human Y437H mutant *MYOC* was inserted in a transcriptionally active genomic locus (H11). A stop cassette prevents the expression of mutant MYOC-DsRed fusion protein until Cre recombinase is introduced. Following Cre expression, the stop cassette is excised allowing the expression of the mutant *MYOC* fused with DsRed only in targeted cells. (**B**) Representative slit lamp images showing that no obvious ocular inflammation is associated with either HAd5-empty or Cre-injected *Tg.CreMYOC^Y^*^437^*^H^* mice (n=6). (**C**) HAd5-Cre was injected intravitreally (2 x10^7^ pfu/eye) in mT/mG fluorescence-based reporter mice and the conversion from tdTomato to GFP was examined 1-week post injection using confocal microscopy. (n=4). (**D-F**) *Tg.CreMYOC^Y^*^437^*^H^* mice were injected intravitreally with HAd5-empty or Cre and MYOC induction was examined in the TM using (**D**) Confocal imaging of DsRed protein in whole mount anterior segment, (**E**) Western blot analysis of various ocular tissues using MYOC antibody showing the presence of human mutant myocilin in anterior segment tissues of Cre-injected eyes; AS-anterior segment; M-empty lane; CS-choroid and sclera, and (**F**) Confocal imaging of DsRed protein in anterior segment cross-section in Cre- and Cre+ *Tg.CreMYOC^Y^*^437^*^H^* mice (n=4). (**G**) RNAscope analysis of *MYOC* (green) and *Myoc* (pink) transcripts in the anterior segment cross-section of Cre- and Cre+ *Tg.CreMYOC^Y^*^437^*^H^*mice. Last panel shows DsRed protein expression in the same slide. Note that DsRed fluorescence may be less compared to other images due to fluorescence quenching during sample processing (TM, trabecular meshwork; CB, ciliary body; SC, Schlemm’s canal). Arrow shows TM.

### Helper Ad5-Cre induces mutant myocilin selectively in the TM

We have previously shown that intravitreal injection of Ad5 exhibits specific tropism to the mouse TM^33–36^. Therefore, we utilized Ad5 expressing Cre recombinase to selectively induce mutant myocilin in the TM of *Tg.CreMYOC^Y^*^437^*^H^* mice (Figure SI-2A,B). Single intravitreal injection of Ad5-empty or Ad5-Cre (2x 10^7^ pfu/eye) was performed in 3-month-old *Tg.CreMYOC^Y^*^437^*^H^* mice. Analysis of anterior segment cross sections from the mouse eyes at 5-weeks post-injection demonstrated robust MYOC-DsRed expression in the TM (Figure SI-2A) of the Ad5-Cre-injected *Tg.CreMYOC^Y^*^437^*^H^* mice. No MYOC-DsRed was detected in the corresponding sections from control mice, nor in any other ocular tissues from the Cre-induced mice, including the retina. Western blot analysis of iridocorneal angle, retina, and sclera clearly demonstrated that mutant myocilin is selectively induced in the TM tissue of 5-weeks Ad5-Cre-injected *Tg.CreMYOC^Y^*^437^*^H^*mice (Figure SI. 2B). Slit-lamp imaging revealed moderate ocular inflammation after Ad5-Cre injections (Figure SI. 2C). To reduce the ocular inflammation, we next utilized helper Ad5 (HAd5), which lacks most of viral sequences except the cis-acting elements essential for viral replication and packaging. HAd5 is known to exhibit minimal immunogenicity while maintaining robust tropism for the tissue of interest^37^. A single intravitreal injection of HAd5 expressing Cre or empty cassette was performed in 3-month-old *Tg.CreMYOC^Y^*^437^*^H^* mice. Slit-lamp imaging demonstrated no visible signs of ocular inflammation in HAd5-empty or HAd5-Cre injected *Tg.CreMYOC^Y^*^437^*^H^*mice (Figure 1B). First, we examined whether HAd5 expressing Cre exhibits functional activity in mouse TM using fluorescence reporter mTmG mice. These mice express tdTomato in all tissue (red fluorescence); expression of Cre induces conversion of tdTomato to GFP (green fluorescence) ^38^. We performed an intravitreal injection of HAd5-empty or HAd5-Cre, and GFP/tdTomato expression was examined using confocal imaging of anterior segment cross-sections (Figure 1C and Figure SI-3). Compared to empty-injected mTmG mice, which only expressed tdTomato, Cre-injected mice exhibited GFP expression selectively in TM cells, and the efficiency of conversion was nearly 90% (Figure SI-3). These data indicate that a single intravitreal injection of HAd5-Cre is highly efficient in transducing the mouse TM.

Next, we investigated whether HAd5-Cre induces mutant myocilin in the TM. A single intravitreal injection of HAd5-Cre or HAd5-empty (2x 10^7^ pfu/eye) was performed in 2-month-old *Tg.CreMYOC^Y^*^437^*^H^* mice and MYOC-DsRed expression in various ocular tissues was evaluated (Figure 1D-F). Whole mount anterior segment imaging demonstrated mutant myocilin expression throughout the TM of Cre-injected *Tg.CreMYOC^Y^*^437^*^H^*mice (Figure 1D). Western blot analysis of anterior segment (AS), retina, and choroid-sclera (CS) tissue lysates demonstrated the presence of MYOC-DsRed protein in the AS, but not in the retina or CS of Cre^+^*Tg.CreMYOC^Y^*^437^*^H^* mice (Figure 1E). MYOC-DsRed protein was detected at ∼75 kDa due to the DsRed tag on mutant MYOC (Figure 1E). We also detected higher molecular weight bands for MYOC, which may represent heteromeric complexes of MYOC, as described previously^39, 40^. Although endogenous myocilin was detected at 50 kDa in all ocular tissues, MYOC-DsRed (75kDa) was only detected in the anterior segment tissues lysates of Cre^+^*Tg.CreMYOC^Y^*^437^*^H^* mice and no mutant MYOC protein was detected in the retina and CS of Cre^+^ *Tg.CreMYOC^Y^*^437^*^H^* mice. Notably, we observed that endogenous myocilin protein was increased in the lysates of iridocorneal angle tissue of Cre-injected *Tg.CreMYOC^Y^*^437^*^H^* mice compared to controls. qPCR analysis of anterior segment tissues using primers specific to mutant MYOC confirmed the presence of MYOC mRNA in Cre-induced *Tg.CreMYOC^Y^*^437^*^H^* mice (Figure SI-4). Confocal imaging of anterior segment cross-sections revealed a robust and selective induction of mutant MYOC in the TM of *Tg.CreMYOC^Y^*^437^*^H^* mice (Figure 1F). Immunostaining for alpha smooth muscle actin (SMA), which predominantly labels the TM and ciliary muscle revealed a strong co-localization of mutant MYOC with SMA in the TM region of Cre-injected *Tg.CreMYOC^Y^*^437^*^H^* mice (Figure SI-5). These data indicate that HAd5-Cre selectively induces mutant MYOC in the TM of *Tg.CreMYOC^Y^*^437^*^H^* mice.

Since mutant *MYOC* is driven by the CAG promoter, which can lead to overexpression of mutant *MYOC*, we next compared mRNA transcript of mutant *MYOC* with endogenous *Myoc* in the TM region using RNA scope (Figure 1G and Figure SI-6). These data also reveal that no mRNA transcript for mutant *MYOC* was detected in Cre^-^*Tg.CreMYOC^Y^*^437^*^H^* mice while abundant endogenous *Myoc* was detected in the TM region. Importantly, Cre^+^ *Tg.CreMYOC^Y^*^437^*^H^* mice displayed the presence of mutant *MYOC* transcript selectively in the TM region. Notably, total transcript measurements in the TM region revealed 5-fold upregulation of endogenous *Myoc* expression in Cre^+^*Tg.CreMYOC^Y^*^437^*^H^* mice compared to Cre^-^*Tg.CreMYOC^Y^*^437^*^H^* mice (Figure SI-6). These data indicates that expression of mutant *MYOC* induces endogenous *Myoc* in the TM. Moreover, the expression of the mutant *MYOC* transcript in Cre-injected eyes is similar to that of the *Myoc* in Cre^-^ *Tg.CreMYOC^Y^*^437^*^H^* mice. Together, these data establish that HAd5-Cre selectively induces mutant myocilin in the TM of *Tg.CreMYOC^Y^*^437^*^H^* mice at a level similar to endogenous *Myoc* without causing ocular inflammation.

### Expression of mutant MYOC reduces TM outflow and elevates IOP significantly in HAd5-Cre-injected *Tg.CreMYOC^Y^*^437^*^H^* mice

Three-month-old *Tg.CreMYOC^Y^*^437^*^H^*mice were injected intravitreally with HAd5-empty or Cre, and IOPs were monitored weekly. Starting from 2-weeks post-injection, Cre-treated mice *Tg.CreMYOC^Y^*^437^*^H^* mice exhibited significantly higher and sustained IOP compared to empty-injected *Tg.CreMYOC^Y^*^437^*^H^* mice (Figure 2A). An independent IOP measurements in conscious mice (without the use of anesthesia) confirmed a pronounced and significant IOP elevation in HAd5-Cre injected *Tg.CreMYOC^Y^*^437^*^H^*mice (Figure SI-7). Measurement of outflow facility using the constant flow infusion method displayed significantly reduced outflow facility at 5-weeks after HAd5-Cre injection in *Tg.CreMYOC^Y^*^437^*^H^* mice compared to the control group (11.75 nL/min/mmHg in Cre^+^ vs. 22.32 nL/min/mmHg in Cre^-^ *Tg.CreMYOC^Y^*^437^*^H^* mice) (Figure 2B). Together, these findings demonstrate that HAd5-Cre induces significant and sustained IOP elevation due to reduced aqueous humor outflow facility from the TM in *Tg.CreMYOC^Y^*^437^*^H^* mice.

**Figure 2:**
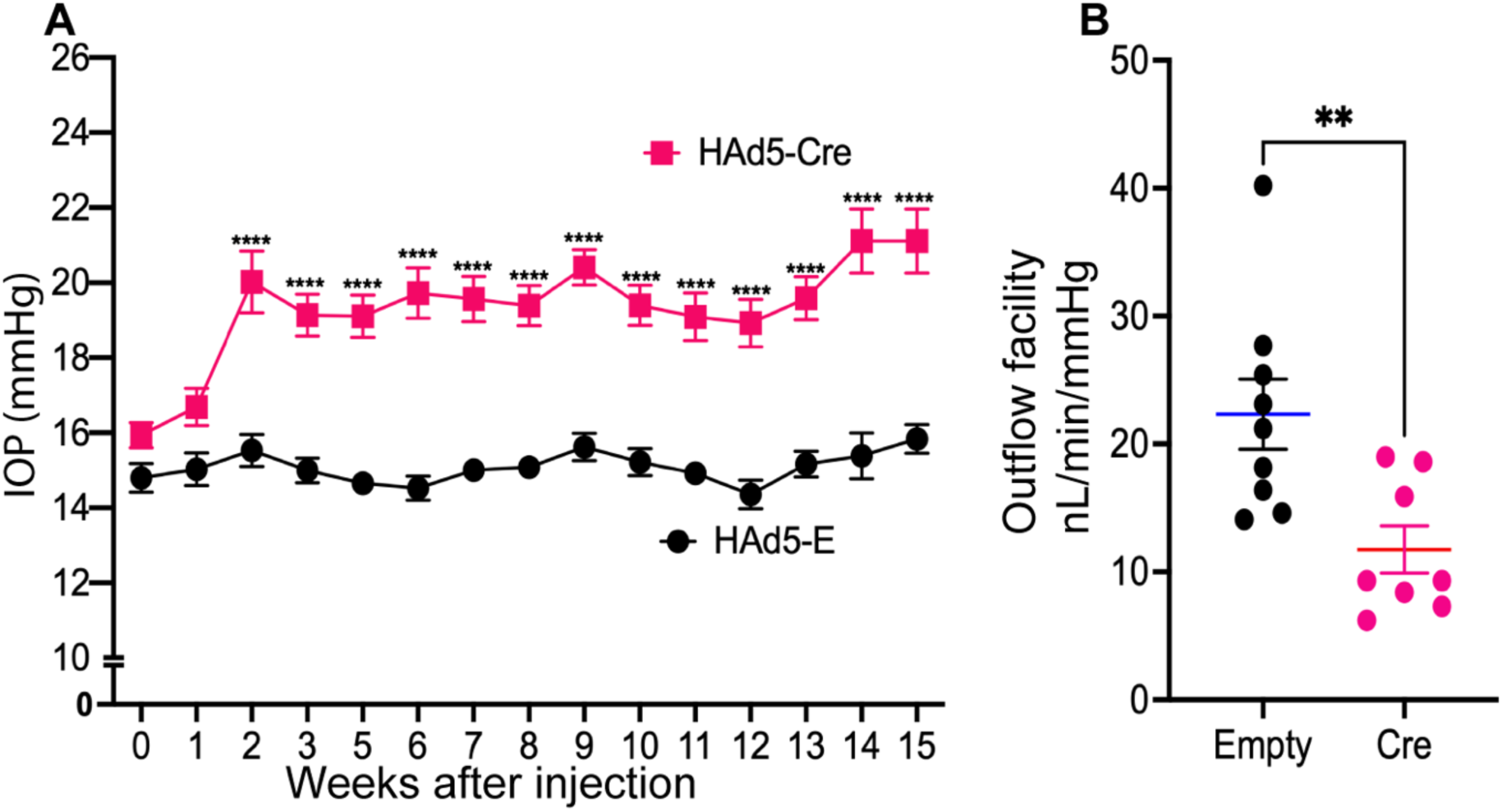
Intravitreal administration of HAd5-Cre elevates IOP and reduces outflow facility in *Tg.CreMYOC^Y^*^437^*^H^*mice. Three-months-old *Tg.CreMYOC^Y^*^437^*^H^*mice received a single intravitreal injection of either HAd5-Empty or Cre in both eyes. (**A**) Weekly IOP measurements demonstrated significant and sustained IOP elevation in Cre-injected *Tg.CreMYOC^Y^*^437^*^H^* mice compared to HAd5-Empty injected mice. (n = 14 in Empty and n=18 in Cre-injected group, analyzed by 2-WAY ANOVA with multiple comparisons, *\*\*\*\*P* <0.0001). (**B**) Outflow facility measurements showed a significant reduction in outflow facility 5-weeks post Cre-injection of *Tg.CreMYOC^Y^*^437^*^H^* mice (n=8) compared to HAd5-Empty-injected mice (n=9) (unpaired t-test, two-tailed, mean ± SEM \*\**P*<0.0072).

### Mutant myocilin induces ultrastructural and biochemical changes in the TM

In POAG, increased outflow resistance is associated with ultrastructural and biochemical changes in the TM including increased extracellular matrix (ECM) deposition, actin cytoskeleton changes and induction of ER stress^11, 15, 41, 42^. We next investigated whether mutant myocilin leads to morphological changes in the TM of *Tg.CreMYOC^Y^*^437^*^H^* mice. *H & E* staining demonstrated open angle and no obvious morphological changes in the anterior chamber structures of Cre^+^*Tg.CreMYOC^Y^*^437^*^H^*mice 5-weeks after injection (Figure SI-8). We next performed transmission electron microscopy (TEM) to examine ultrastructural changes in the TM. Low magnification TEM analysis demonstrated that iridocorneal angle is open in both empty and Cre-injected *Tg.CreMYOC^Y^*^437^*^H^* mice (Figure 3A). Higher magnification TEM images revealed loosely bound collagen fibers, ECM deposition, and disrupted TM integrity in Cre^+^ *Tg.CreMYOC^Y^*^437^*^H^* mice compared to Cre^-^ *Tg.CreMYOC^Y^*^437^*^H^*mice (Figure 3B). We further confirmed these findings using immunostaining. Anterior segments were immunostained with antibodies for fibronectin (FN) and actin (Figure SI-9A). Both FN and actin were significantly increased in the TM region of Cre-injected *Tg.CreMYOC^Y^*^437^*^H^*mice (Figure SI-9B). Previous studies have shown that mutant myocilin expression in TM induces ER stress^30, 43–46^. Therefore, we examined whether mutant myocilin expression induces ER stress markers in the TM of Cre-induced *Tg.CreMYOC^Y^*^437^*^H^* mice (Figure 3C). Immunostaining (Figure SI-10) and Western blot analysis (Figure 3C) demonstrated significantly increased ER stress markers including GRP78, ATF4 and CHOP selectively in the TM of Cre-injected *Tg.CreMYOC^Y^*^437^*^H^*mice. Together, these findings indicate that expression of mutant myocilin induces ultrastructural and biochemical changes in the TM leading to its dysfunction and IOP elevation in *Tg.CreMYOC^Y^*^437^*^H^* mice.

**Figure 3.**
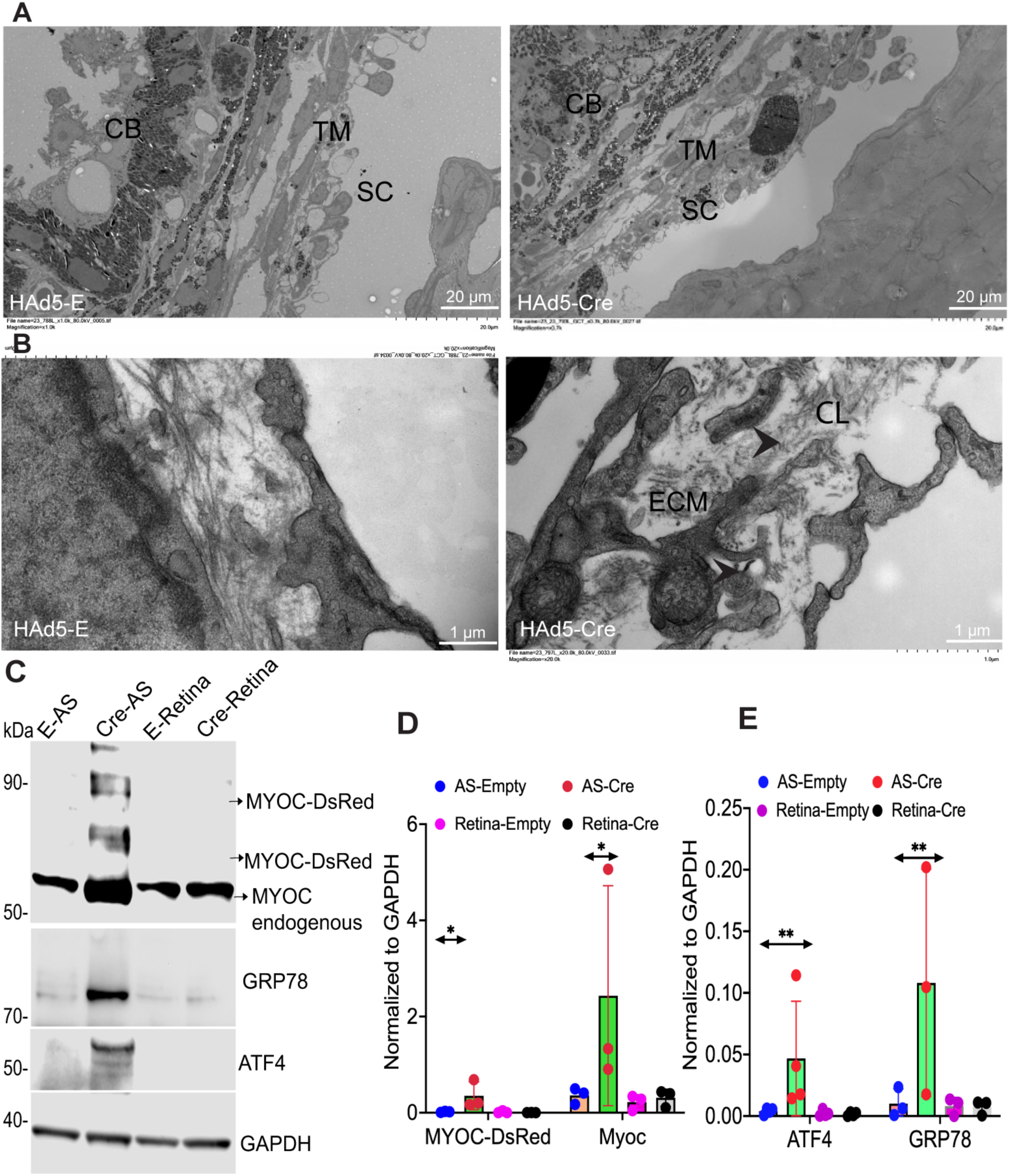
Mutant MYOC-induced ocular hypertension is associated with ultrastructural and biochemical changes in the TM. Representative low. **(A)** and high (**B**) magnification TEM images of *Tg.CreMYOC^Y^*^437^*^H^* mice 8-weeks post HAd5-Empty and HAd5-Cre injection showing the presence of loosely bound collagen fibers, ECM deposition and loss of TM integrity in juxtacanalicular connective tissue (JCT) region of Cre-injected *Tg.CreMYOC^Y^*^437^*^H^* mice (n=4 in each group). (TM, trabecular meshwork; CB, ciliary body; SC, Schlemm’s canal; CL, Collagen fibers; ECM, Extra cellular matrix). (**C-D**) Western blot and densitometric analyses showing that mutant MYOC induces ER stress in the anterior segment tissue lysates of Cre-injected *Tg.CreMYOC^Y^*^437^*^H^* mice. (n=3). E-HAd5-empty; AS-anterior segment (2-WAY ANOVA with multiple comparisons (*\*\*P*=0.0053 & *\*P*=0.0216).

### Mutant MYOC-induced sustained IOP elevation leads to functional and structural loss of RGCs

We next evaluated whether sustained IOP elevation induced by mutant MYOC is sufficient to cause functional and structural loss of RGCs. To evaluate functional loss of RGCs, we performed PERG at 5, 10, and 15 weeks after injection (Figure 4A-C). Representative PERG graphs and their analysis demonstrated no significant effect on PERG at 5-weeks, but significantly reduced PERG amplitudes and increased latencies were observed at 10 and 15 weeks Cre-injected *Tg.CreMYOC^Y^*^437^*^H^*mice (Figure 4A-C). To determine the structural loss of RGCs, we next performed whole mount retina staining with RBPMS antibody (Figure 4D-E). As shown in representative RBPMS images, Cre-injected *Tg.CreMYOC^Y^*^437^*^H^* mice exhibited moderate loss of RGCs in the peripheral retina after 15 weeks of injection (Figure 4E). RGC counting further confirmed a significantly reduced RGCs in the periphery of Cre-injected *Tg.CreMYOC^Y^*^437^*^H^*mice compared to controls at 15 weeks of injection (Figure 4E). Overall, there was a 33% loss of RGCs in the peripheral retina of Cre-injected *Tg.CreMYOC^Y^*^437^*^H^* mice compared to controls. We did not observe RGC loss at 10-weeks after Cre-injection in *Tg.CreMYOC^Y^*^437^*^H^* mice (Figure SI-11). These data indicate that sustained IOP elevation induced by mutant myocilin leads to RGC functional and structural loss of RGCs in *Tg.CreMYOC^Y^*^437^*^H^* mice.

**Figure 4:**
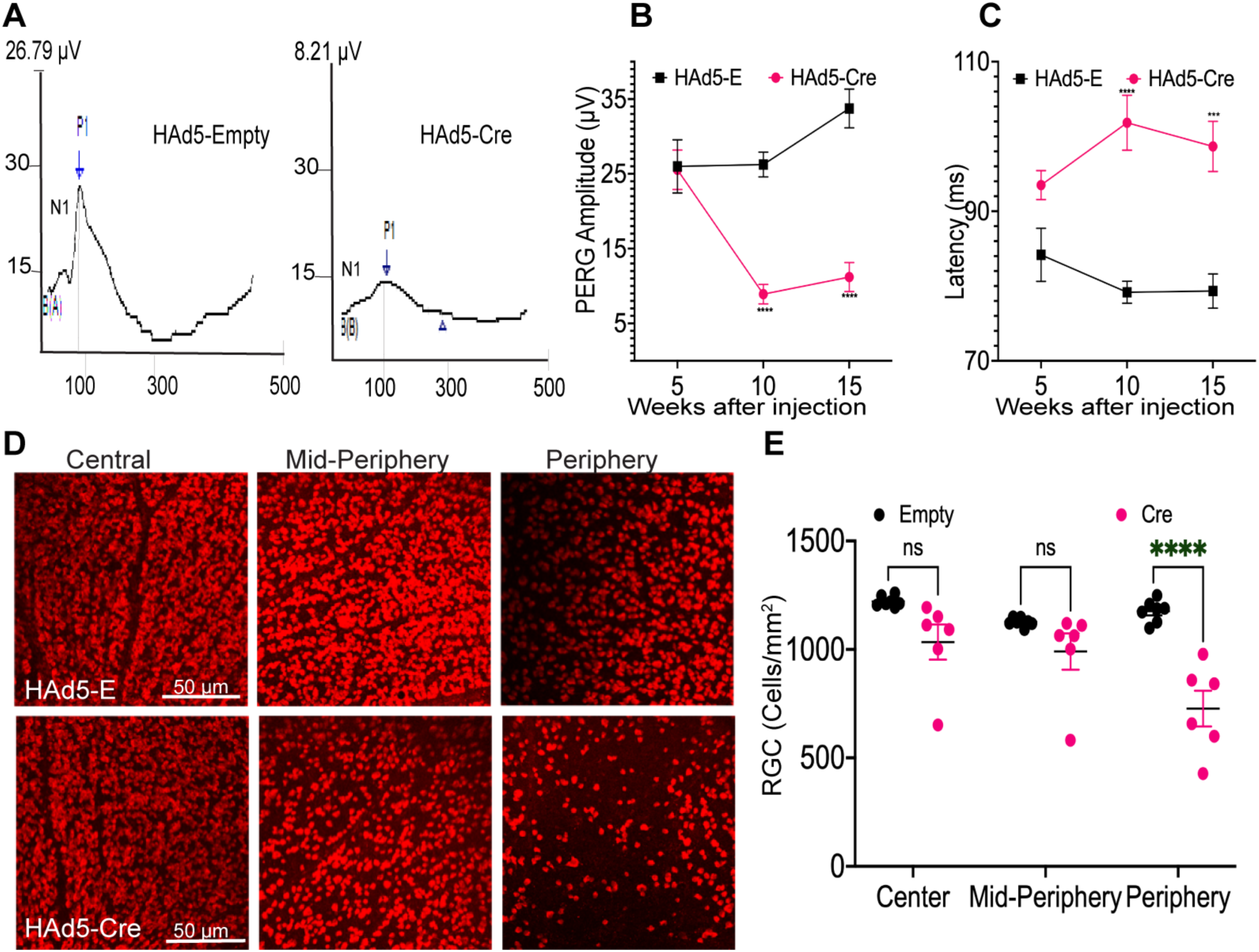
Sustained IOP elevation leads to functional and structural loss of RGCs in Cre-injected *Tg.CreMYOC^Y^*^437^*^H^* mice: 3 to 6-months-old *Tg.CreMYOC^Y^*^437^*^H^* mice were injected intravitreally with HAd5-Empty or HAd5-Cre in both eyes, and IOP was monitored weekly to ensure IOP elevation. PERG was performed at 5-, 10- and 15-weeks post treatment to assess the function of RGCs. A representative PERG graph (**A**) and its analysis (**B-C**) demonstrated significantly reduced PERG amplitude (**B**) and increased latency (**C**) starting from 10-weeks post Cre injection indicating functional loss of RGCs in Cre^+^*Tg.CreMYOC^Y^*^437^*^H^* mice. (n = 7 in HAd5-Empty and n=6 in HAd5-Cre), 2-WAY ANOVA with multiple comparisons (\*\*\*\**P* <0.0001). (**D-E**) RGCs loss was further analyzed using whole mount retina staining with RBPMS antibody. Representative image of RBPMS staining of different regions of retina (**D**) and its analyses (**E**) revealed a significant loss (33%) of RGCs in 15-weeks post injection of HAd5-Cre in *Tg.CreMYOC^Y^*^437^*^H^* mice compared to HAd5-Empty injected mice. (n = 7 HAd5-Empty and 6 in HAd5-Cre), 2-WAY ANOVA with multiple comparisons (\*\*\*\**P*<0.00001).

### Mutant MYOC-induced sustained IOP elevation leads to optic nerve degeneration in *Tg.CreMYOC^Y^*^437^*^H^* mice

We investigated whether sustained IOP elevation induced by mutant myocilin leads to optic nerve degeneration, using PPD staining (Figure 5A). Representative images of PPD stained ON from Cre-injected *Tg.CreMYOC^Y^*^437^*^H^*mice revealed optic nerve degeneration as evident from darkly stained axons, active gliosis, and glial scar formation. Approximately 20% and 45% axonal loss was observed in Cre-injected mice at 10 and 15-weeks respectively compared to empty-injected *Tg.CreMYOC^Y^*^437^*^H^* mice (Figure 5B). To further confirm these findings, we next performed immunostaining for GFAP on retinal cross-sections from Cre-injected *Tg.CreMYOC^Y^*^437^*^H^* mice (Figure SI-12). Immunostaining for GFAP revealed a prominent increase in GFAP reactivity in the ONH region suggesting axonal injury. Axonal degeneration was associated with decreased neuronal marker, Tuj1 and increased mitochondrial accumulation (increased TOM20) in the ONH region (Figure SI-12). These data establish that sustained IOP elevation induces ON degeneration and gliosis in the ONH of Cre-injected *Tg.CreMYOC^Y^*^437^*^H^* mice.

**Figure 5:**
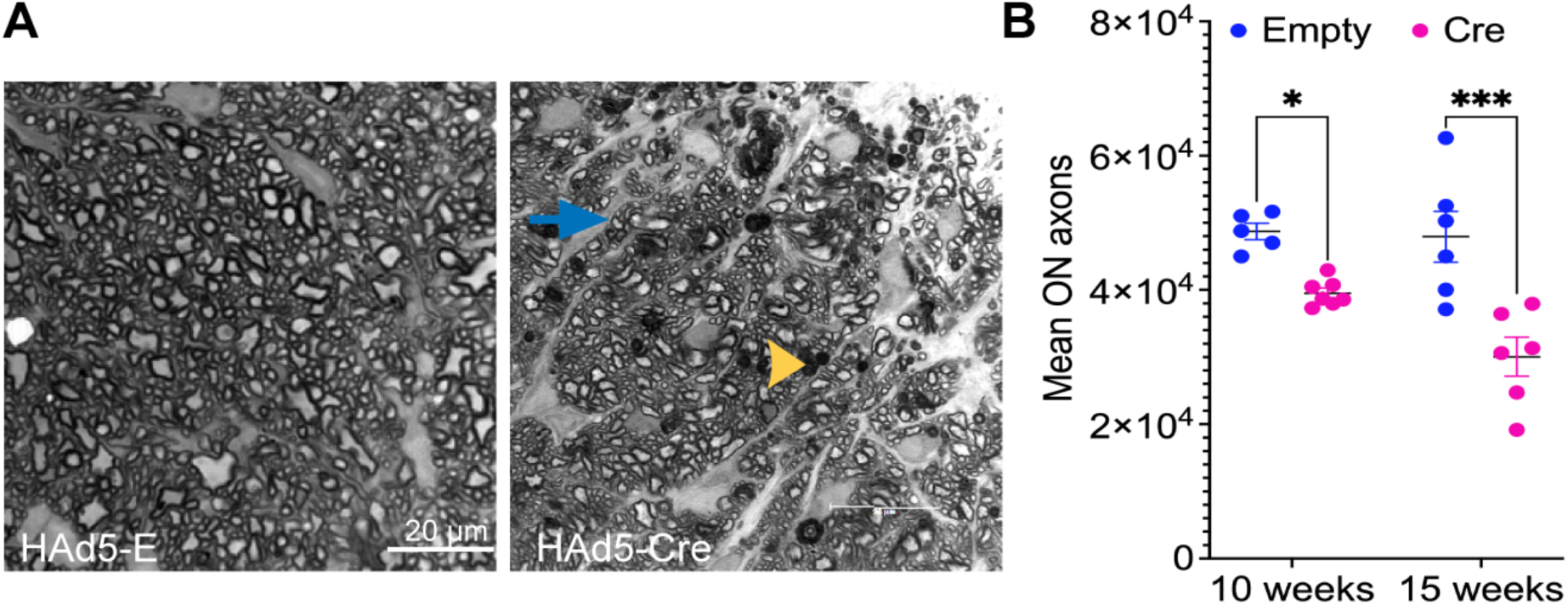
Sustained IOP elevation leads to optic nerve degeneration in Cre-injected *Tg.CreMYOC^Y^*^437^*^H^* mice. Optic nerves were subjected to PPD staining to assess optic nerve degeneration. (**A**) Representative images of PPD-stained optic nerves showing mild axonal degeneration as evident from darkly stained axons (yellow arrow) and the presence of glial scar formation (blue arrow) in Cre-injected *Tg.CreMYOC^Y^*^437^*^H^* mice and (**B**) the mean axonal counts showing a significant loss of ON axons (20% at 10-weeks and 45% at 15-weeks post injection in Cre-induced *Tg.CreMYOC^Y^*^437^*^H^* mice. (n = 5 in Empty and n = 7 at 10-weeks post injection, and n=6 in Empty and n=6 at 15-weeks post injection, unpaired t-test, two-tailed, \*\*\*\**P* < 0.0001).

### Impaired axonal transport at the ONH precedes neuronal loss in *Tg.CreMYOC^Y^*^437^*^H^*mice

Since *Tg.CreMYOC^Y^*^437^*^H^* mice exhibit well-defined timelines for RGC axon loss, we further sought to understand the early pathogenic events preceding neuronal loss. RGC loss and ON degeneration studies suggested that axonal changes at the ONH may be the first site of damage in ocular hypertensive *Tg.CreMYOC^Y^*^437^*^H^* mice. We therefore hypothesize that IOP-induced neurodegenerative changes in the ONH precede neuronal loss. Since *Tg.CreMYOC^Y^*^437^*^H^* mice did not show significant structural loss of RGCs at 10-weeks after Cre-injection (Figure SI-11), we examined whether axonal dysfunction occurs at the ONH prior to RGC loss at 7-weeks after Cre-injection. To test this, 15-month-old *Tg.CreMYOC^Y^*^437^*^H^*mice were injected intracamerally with HAd5-empty or Cre and IOPs were monitored. IOP measurements confirmed a significant IOP elevation at 6-weeks after Cre-injection of *Tg.CreMYOC^Y^*^437^*^H^* mice (Figure 6A). PERG, which measures the function of RGC soma revealed no significant functional loss of RGC soma at 5-weeks after Cre-injection of *Tg.CreMYOC^Y^*^437^*^H^* mice (Figure 4A-C). Whole mount retinal staining with RBPMS revealed no significant loss of RGCs at 10-weeks after Cre-injection of *Tg.CreMYOC^Y^*^437^*^H^* mice (Figure SI-11). Notably, VEP, which measures post-retinal function of the visual system including ON and visual centers of the brain demonstrated significant loss of VEP amplitudes at 7-weeks suggesting axonal dysfunction in Cre^+^*Tg.CreMYOC^Y^*^437^*^H^* mice (Figure 6B). To understand whether axonal dysfunction occurs at early stages of neuronal loss due to sustained IOP, we examined anterograde transport of fluorescently labeled CTB as described previously^47^. At 7-weeks post injection, HAd5-empty or HAd5-Cre-injected *Tg.CreMYOC^Y^*^437^*^H^* mice were intravitreally injected with green fluorescently tagged CTB dye (Figure 6C). 48-hours after injections, anterograde transport of CTB through the optic nerve and superior colliculus (SC) was monitored via fluorescence microscopy. As expected, CTB was transported to the SC in HAd5-empty injected *Tg.CreMYOC^Y^*^437^*^H^* mice. However, CTB transport was completely blocked at the ONH and no CTB was observed in the SC of Cre-injected *Tg.CreMYOC^Y^*^437^*^H^*mice. CTB fluorescence measurement in SC demonstrated a significant (∼72%) loss of CTB in SC of Cre-injected *Tg.CreMYOC^Y^*^437^*^H^* mice (Figure 6D). Previous studies have shown that axonal transport deficits are age dependent. Since we observed significant axonal transport in 15-month-old Cre^+^*Tg.CreMYOC^Y^*^437^*^H^*mice, we next explored whether axonal transport deficits are observed in younger Cre^+^*Tg.CreMYOC^Y^*^437^*^H^*mice. 4-month-old *Tg.CreMYOC^Y^*^437^*^H^*mice were intravitreally injected with HAd5-empty or HAd5-Cre; at 7-weeks post-injection, ocular hypotensive eyes were analyzed for CTB transport (Figure SI-13A,C). Similar to 15-months old mice, CTB transport was blocked at the ONH and a significant reduction (65%) of CTB transportation to the SC was observed in 4-month-old *Tg.CreMYOC^Y^*^437^*^H^* mice (Figure SI-13B,C). Altogether, these data indicate that ocular hypertension impairs axonal transport blockage at the ONH prior to RGC loss in Cre^+^*Tg.CreMYOC^Y^*^437^*^H^*mice in an age-independent manner.

**Figure 6:**
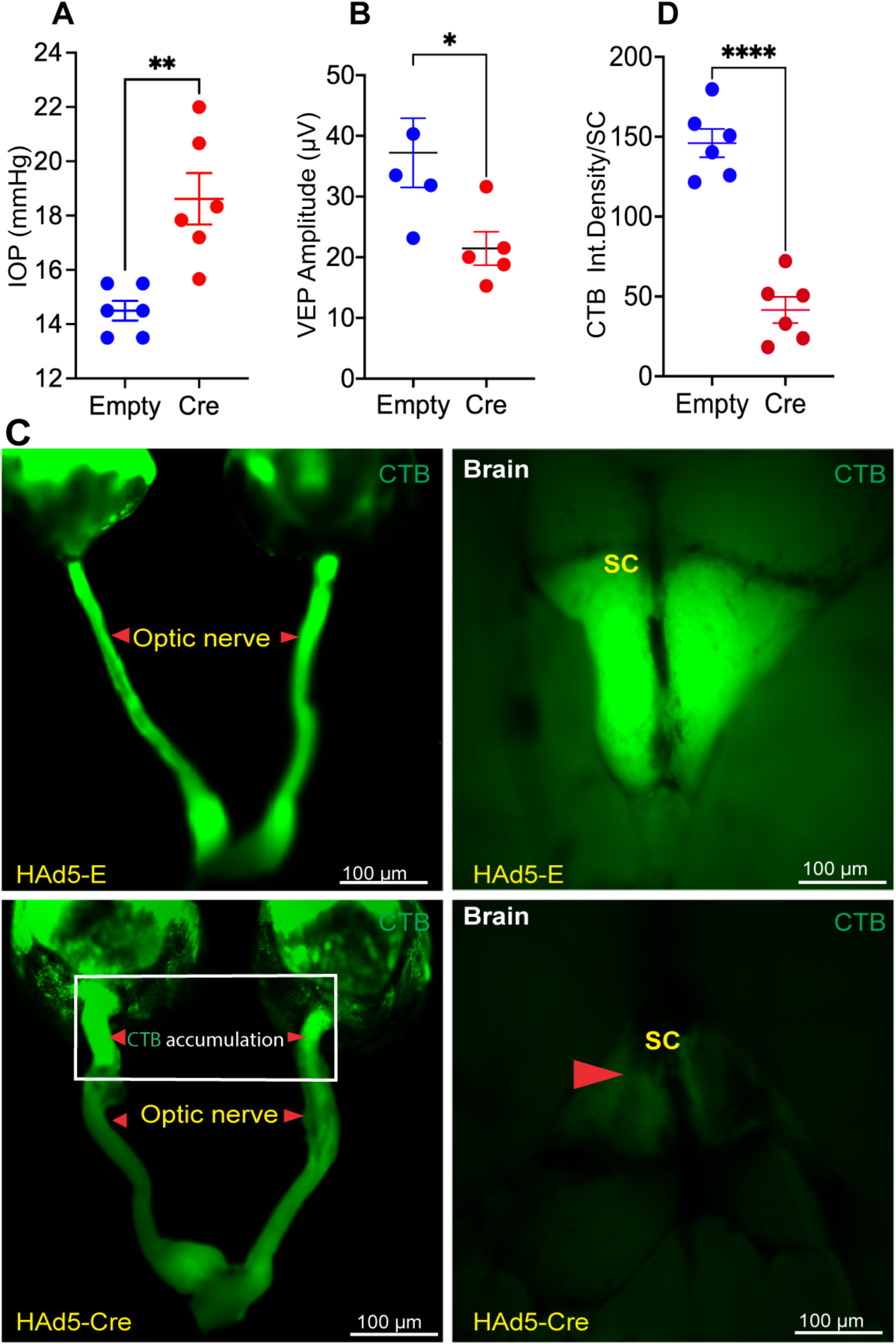
Anterograde transport deficits precede RGC degeneration in *Tg.CreMYOC^Y^*^437^*^H^* mice: 15-months-old *Tg.CreMYOC^Y^*^437^*^H^*mice were injected intravitreally with HAd5-Empty or Cre and glaucoma phenotypes and anterograde axonal transport mechanisms were investigated. **(A)** IOP measurement revealed sustained and significant IOP elevation at 6-weeks post injection (n=6; unpaired t-test; *P*=0.0058). **(B)** VEP measurements demonstrated a significant loss of post-retinal visual pathway function in 7-weeks post Cre injection in *Tg.CreMYOC^Y^*^437^*^H^* mice (n=5; unpaired t-test; *P*=0.0374). (**C-D**) Mutant MYOC-induced ocular hypertension leads to axonal transport deficits in Cre-injected *Tg.CreMYOC^Y^*^437^*^H^* mice. Representative images of CTB fluorescence (**C**) and its analysis (**D**) in 7-weeks post HAd5-Empty or Cre injection in *Tg.CreMYOC^Y^*^437^*^H^* mice. Empty-injected *Tg.CreMYOC^Y^*^437^*^H^* mice exhibited an uninterrupted transport of CTB along the entire length of optic nerve to the SC. However, CTB transport was blocked significantly at the ONH region and no CTB was detected in SC of 7-weeks post Cre-injection in *Tg.CreMYOC^Y^*^437^*^H^*mice (n = 6 in each group).

### Ultrastructural changes in RGC axons are associated with impaired axonal transport in ocular hypertensive Cre^+^*Tg.CreMYOC^Y^*^437^*^H^*mice

To further gain molecular insights about impaired axonal transport, we analyzed CTB transport in cross-section of retina, along with the ON. CTB transport was blocked in the proximal region of the ON in Cre-injected *Tg.CreMYOC^Y^*^437^*^H^* mice (Figure 7A). TEM analysis of the ON at 8-weeks after Cre-injection demonstrated ultrastructural changes including loss of neurofilament and microtubule structures (Figure 7B-C). The presence of myelin around axons in Cre-injected mice indicated intact axons, but accumulation of intracellular materials and loss of microtubules and neurofilaments suggest that these ultrastructural changes in ON axons precede axonal degeneration. To confirm the TEM findings, we performed immunostaining on retinal cross-sections obtained from Cre^-^ and Cre^+^ *Tg.CreMYOC^Y^*^437^*^H^* mice 7-weeks after injection (Figure SI-14). Cre^+^*Tg.CreMYOC^Y^*^437^*^H^* mice exhibited a dramatic increase in GFAP reactivity, decreased microtubules (Tuj1) and loss of neurofilament (NF) loss in the ONH region compared to Cre^-^*Tg.CreMYOC^Y^*^437^*^H^* mice. Importantly, we observed increased mitochondrial accumulation in RGC soma (Figure SI-14) and axons at the ONH region in Cre^+^*Tg.CreMYOC^Y^*^437^*^H^* mice after 12-weeks of injection (Figure SI-12). Together, our studies suggest that loss of microtubules and neurofilaments, which are required for axonal transport, is associated with impaired axonal transport leading to accumulation of defective organelles and causing axonal dysfunction.

**Figure 7:**
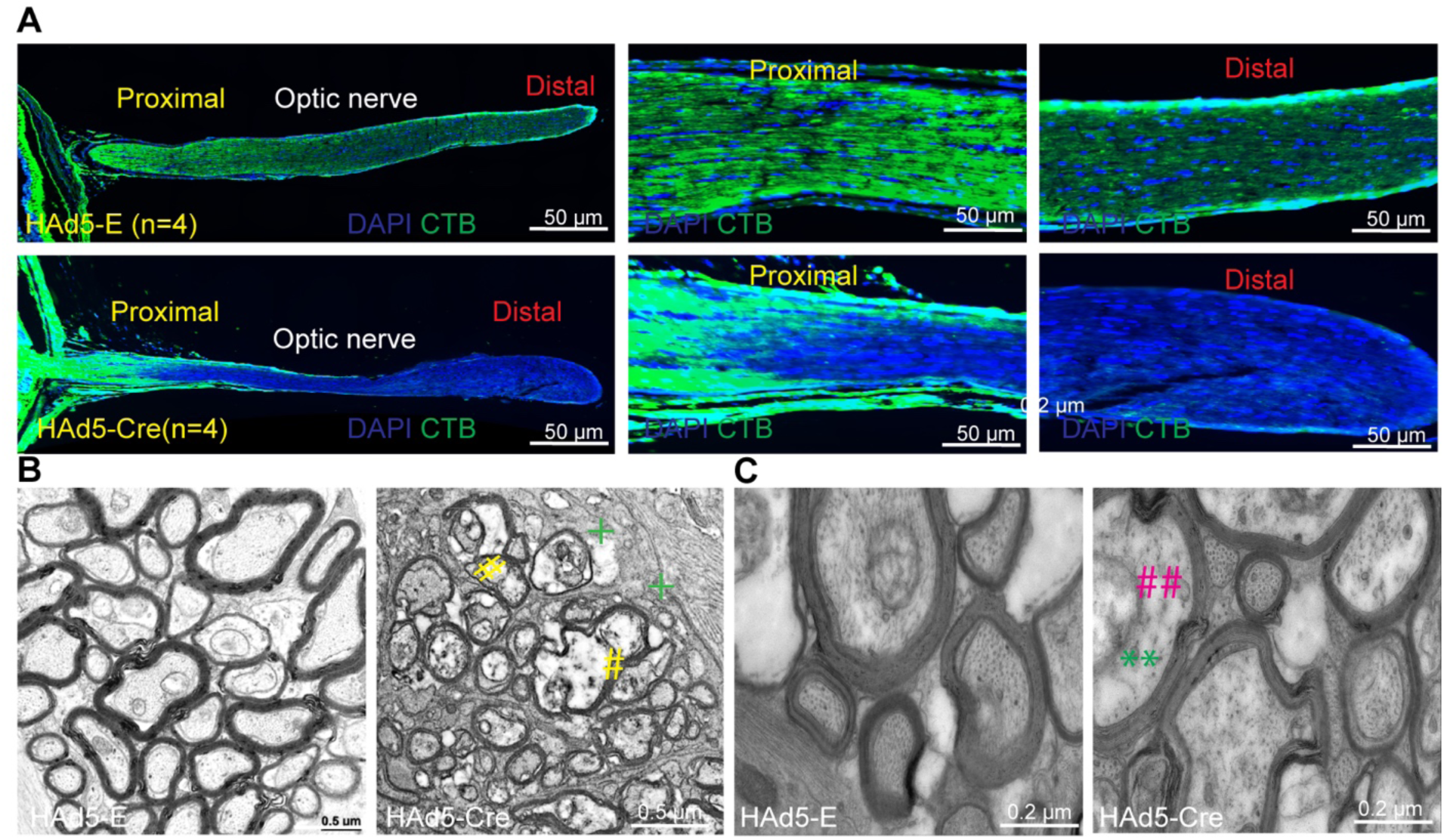
Reduced microtubules and neurofilament are associated with axonal transport deficits in the ONH region. (**A**) Whole eyes along with ON from above CTB-injected *Tg.CreMYOC^Y^*^437^*^H^* mice were sectioned to image CTB transport along ON. Retinal cross-sections demonstrated the blockage of CTB transport in the proximal region of ON in Cre-injected *Tg.CreMYOC^Y^*^437^*^H^* mice. (n=3). (**B**) Low and (**C**) high magnification TEM analysis of optic nerves Empty or Cre-injected *Tg.CreMYOC^Y^*^437^*^H^* mice, which demonstrated cytoskeleton degeneration, organelle accumulation and the presence of glial scar prior to RGC degeneration in 7-weeks post Cre injection of *Tg.CreMYOC^Y^*^437^*^H^* mice (n=4). (organelle accumulation; + glial scar). (**Neurofilament; ## microtubule).

### Axonal transport remains intact in ocular hypertensive mice injected with microbeads despite significant RGC loss

To determine whether impaired axonal transport is a unique and early pathological hallmark of Cre^+^*Tg.CreMYOC^Y^*^437^*^H^* mice, we next compared axonal transport mechanisms in a mouse model of microbead (MB)-induced OHT. MB-induced OHT model is widely utilized model to study glaucomatous neurodegeneration^21, 48^. 4-month-old C57BL/6J mice were intracamerally injected with magnetic MBs, which block TM outflow, elevating IOP as described previously^49, 50^. Weekly IOP measurements demonstrated that about 60% of MB-injected eyes exhibited significant IOP elevation (≥ 4mmHg) (Figure 8A). The eyes that do not show IOP elevation likely reflect technical failure to keep the MBs in the outflow pathway. The ocular hypertensive eyes (IOP elevation > 4mmHg) at 6-weeks post MB-injection were selected for RGC functional and structural analysis, using PERG measurements and RBPMS staining respectively. PERG (Figure 8B) and RBPMS staining (Figure SI-15 and Figure 8C) demonstrated significant structural and functional loss of RGCs in the MB-injected eyes. GFAP staining (Figure SI-15) demonstrated increased astrocyte labeling is associated with RGC loss in MB-injected mice. We next investigated whether ocular hypertensive MB-injected mice exhibit axonal transport deficits as seen in Cre^+^*Tg.CreMYOC^Y^*^437^*^H^* mice. 4-month-old C57 mice were injected with PBS or MB and IOPs were monitored (Figure SI-16A). PERG measurements in ocular hypertensive eyes at 3 weeks post MB-injection revealed a significant functional loss of RGCs in MB-induced OHT mice (Figure SI-16B). CTB was injected intravitreally at week 4 after MB injection and CTB transport to the SC was examined (Figure 8D-E). Despite significant functional loss of RGCs, we observed that CTB transport to the SC was completely intact (Figure 8D-E). These data indicates that axonal transport persists despite significant RGC degeneration in a mouse model of MB-induced OHT.

**Figure 8:**
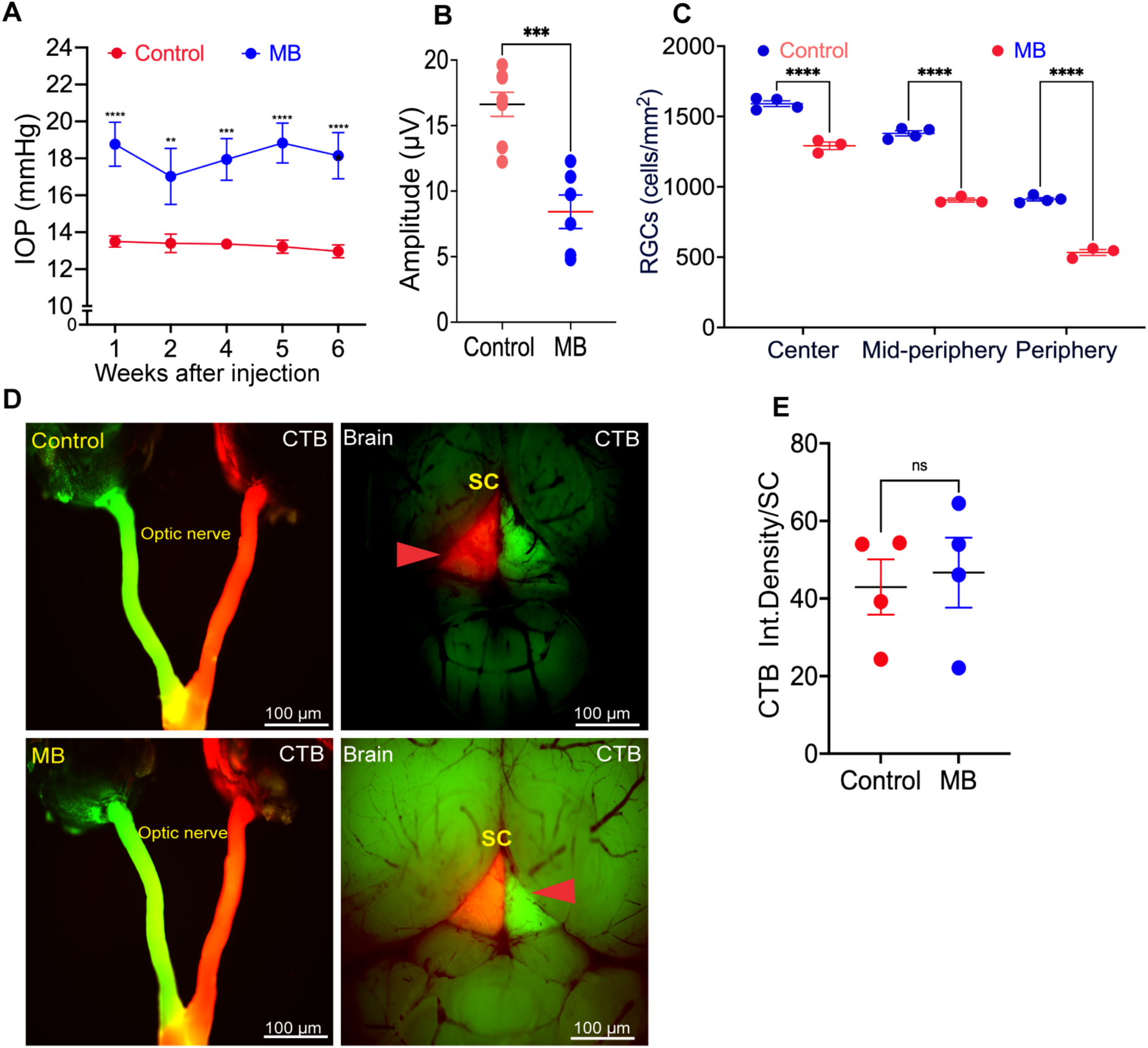
Anterograde axonal transport remains intact despite RGC degeneration in ocular hypertensive mice injected with microbeads: 4-month-old C57 mice were injected intracamerally with PBS or microbeads (MB). Glaucoma phenotypes and anterograde axonal transport mechanisms were investigated. **(A)** IOP measurement revealed sustained and significant IOP elevation in MB-injected mice starting from the first week of injection. Note that only 50% of eyes injected with MB elevated IOP more than 4 mmHg. The graph includes eyes that showed sustained IOP elevation of 4mmHg or more than4mmHg. (n=10 for control, n=7 MB; analyzed by 2-WAY ANOVA with multiple comparisons, \*\*\*\**P* <0.0001). **(B)** PERG measurements demonstrated a significant functional loss of RGCs 6-weeks post MB injection (n=4 control, n=4 MB; (unpaired t-test, two-tailed, mean ± SEM \*\*\**P*=0.0002). (**C**) RGCs analysis from whole mount RBPMS staining of retina revealed a significant loss of RGCs 6-weeks post MB-injection (n=4 for control, n=3 MB; analyzed by 2-WAY ANOVA with multiple comparisons, \*\*\*\**P* <0.0001). **(D)** Representative images of CTB fluorescence in the optic nerve and SC and (**E**) its analysis in ocular hypertensive mice 4-weeks post MB injection. CTB transportation remained intact despite significant RGC loss in MB-injected mice. (n = 4 in each group; unpaired t-test; ns=not-significant).

## Discussion

In the present study, we describe the development of novel Cre-inducible mouse model of MYOC-associated POAG. We report that HAd5 drives the TM-specific expression of mutant MYOC leading to TM dysfunction and IOP elevation. Importantly, a single intravitreal injection of HAd5-Cre is sufficient to induce sustained IOP elevation, which leads to glaucomatous neurodegeneration in a highly predictable manner. Induction of mutant MYOC via Cre expression in adult mice allowed us to study early events of glaucomatous damage to the TM and IOP-induced neurodegeneration. Importantly, we show that sustained IOP elevation impairs axonal transport at the ONH prior to RGC soma and axonal loss. These mechanistic insights into early events of TM dysfunction and axonal degeneration provide valuable targets to develop IOP-lowering and neuroprotective therapies for glaucoma.

Previously, we have developed a transgenic mouse model (*Tg-MYOC^Y^*^437^*^H^*) by random insertion of mutant MYOC^30^. This transgenic mouse model exhibited robust glaucoma phenotypes on mixed background, and we discovered the pathological role of chronic ER stress in the pathophysiology of glaucomatous TM damage using this model^29, 30, 33^. However, these mice exhibited mild glaucoma phenotypes with subsequent breeding onto pure C57BL/6J background ^36, 51, 52^. We reasoned this to silencing of transgene likely due to the toxic nature of mutant MYOC. In addition, our previous transgenic mouse model presented several limitations including lack of robust glaucomatous neurodegeneration on pure C57 background, unknown copies of mutant MYOC and lack of specific antibodies to distinguish human versus mouse myocilin. Development of *Tg.CreMYOC^Y^*^437^*^H^*mice has overcome these key limitations. These include targeted insertion of mutant MYOC in the preselected intergenic and transcriptionally active genomic locus (H11). This allows stable and site-specific transgene integration of single copy of mutant MYOC. Also, *Tg.CreMYOC^Y^*^437^*^H^* mice are developed on pure C57BL/6J background. DsRed is fused with mutant *MYOC*, which allowed us to distinguish mutant *MYOC* from endogenous *Myoc*. Indeed, RNA scope analysis demonstrated the expression levels of mutant *MYOC* is similar to endogenous *Myoc*. This was further supported by Western blot analysis, which demonstrated protein levels of human mutant MYOC fused with DsRed is comparable to endogenous MYOC in Cre^-^*Tg.CreMYOC^Y^*^437^*^H^* mice. These data indicates that glaucoma phenotypes in *Tg.CreMYOC^Y^*^437^*^H^* mice are not a result of overexpression of mutant MYOC. Both RNA scope and Western blot clearly demonstrated that expression of mutant *MYOC* induced endogenous myocilin gene and protein respectively. This is consistent with previous findings that TM cells induce MYOC in response to various insults including glucocorticoid treatment ^27, 53, 54^. Although the exact function of endogenous myocilin is unclear, previous studies have suggested its role as a stress-response pathway ^55^. It is likely that misfolded mutant MYOC interacts with endogenous myocilin protein causing its accumulation in the ER^39^, further contributing to the TM dysfunction.

The Cre-*LoxP* recombinase system is an effective and widely used experimental tool to investigate gene of interest in tissue/cell and/or time-specific manner^56^. Cre-inducible mice are often crossed with tissue-specific Cre mouse lines to induce Cre-expression. In our study, we employed viral vector to induce Cre expression in TM. Crossing *Tg.CreMYOC^Y^*^437^*^H^* mice with TM-specific Cre mouse line would be ideal to induce mutant MYOC in TM. However, there are no TM-specific Cre mouse lines available. The global Cre mouse lines would express mutant MYOC in other cell types including in ciliary body and retina, which can directly result in cell death of ciliary epithelium or RGCs. Previous studies have shown that mutant MYOC is toxic to cells ^30^ and degenerations of the non-spigmented epithelium in the ciliary body was observed in older *Tg-MYOC^Y^*^437^*^H^*mice ^52^. We also observed that induction of mutant MYOC selectively in RGCs via an intravitreal injection of AAV2-Cre leads to severe RGC loss within 3 weeks of injection in *Tg.CreMYOC^Y^*^437^*^H^* mice (Data not shown). A recent study has shown that Matrix Gla (MGP)-Cre mouse line exhibits Cre activity in TM and other ocular cells relevant to glaucoma pathology^57^. Our future studies will be directed towards using MGP-Cre mouse line to induce mutant MYOC in TM and other ocular cells that are relevant to glaucoma pathology.

Our studies show that *Tg.CreMYOC^Y^*^437^*^H^* mice exhibit all features of human POAG including IOP elevation and reduced outflow facility due to TM dysfunction. Like POAG, we observed ultrastructural and biochemical changes associated with TM dysfunction as evident from increased ECM deposition, actin and ER stress in Cre-injected mice. Importantly, sustained IOP elevation leads to progressive axonal degeneration, which is associated with RGC loss. Contrary to other existing glaucoma models, most of *Tg.CreMYOC^Y^*^437^*^H^* mice develop ocular hypertension and exhibit neuronal loss. Nearly all eyes treated with Cre developed IOP elevation and glaucomatous neurodegeneration. Glaucoma phenotypes in Cre-inducible *Tg.CreMYOC^Y^*^437^*^H^* mice are highly predictable, and the exact timeline of each phenotype can be easily tracked. Mutant MYOC is induced within a week, elevating IOP starting from 2-weeks of Cre injection, which is followed by axonal transport deficits at 7 weeks. Axonal loss is observed at 10-weeks and RGC soma loss is observed at 15-weeks. These features make *Tg.CreMYOC^Y^*^437^*^H^* mouse model highly attractive to study both TM dysfunction and neuronal loss in glaucoma. Although we did not investigate long-term effects of mutant MYOC on TM dysfunction and neuronal loss, it is likely that IOP elevation will sustain throughout the life of mice resulting into severe neuronal loss as observed in POAG.

Our study further demonstrates that sustained IOP elevation impairs axonal transport in the ONH prior to axonal degeneration and RGC structural loss. Axonal transport deficits are considered an important pathological feature of glaucomatous neurodegeneration^58, 59^. Previous studies have shown that impaired axonal transport is associated with glaucomatous neurodegeneration ^5, 49, 60–62^. We have recently demonstrated that axonal transport persists during initial stages of axonal degeneration in mouse model of glucocorticoid-induced glaucoma^47^. In addition, several other studies have shown that impaired axonal transport in ONH region occurs in an inducible model of ocular hypertension^4–6, 47, 63^. However, it is poorly understood impaired axonal transport is a result or cause of RGC degeneration. In 7-weeks of Cre-injected *Tg.CreMYOC^Y^*^437^*^H^* mice, which demonstrated complete blockage of axonal transport, we observed no significant loss of axon as evident from intact myelin. However, TEM analysis demonstrated intracellular ultrastructural changes including swollen axons, increased mitochondrial accumulation and gliosis indicating axonal dysfunction. Interestingly, we observed a significant loss of VEP, which measures post-retinal visual pathways (ON and visual center of the brain)^64^ while RGC function (measured by PERG) was not significantly affected until 10-weeks of injection. Together, these findings indicate that impaired axonal transport at the ONH precedes RGC soma and axonal loss. Since axonal transport is essential for RGC soma survival including transport of organelles, synaptic components, vesicles, and neurotrophic factors, it is likely that impaired axonal transport leads to RGC axonal degeneration and soma loss. Consistent with this, we observed increased mitochondrial accumulation in RGC soma and ON axons in 12-weeks Cre-injected *Tg.CreMYOC^Y^*^437^*^H^* mice.

It is interesting to note that axonal transport persisted despite significant RGC loss in MB-model. A previous study has shown that significant axonal transport is observed in MB model initially at the SC, which then progresses proximally to optic nerve^49^. Moreover, axonal transport deficits were only observed in aged animals despite both young and old mice show RGC loss. This suggest that impaired axonal transport is unlikely to cause RGC loss. Impaired axonal transport in *Tg.CreMYOC^Y^*^437^*^H^*mice occurred at early stage of neurodegeneration, and it was observed in both young and old mice. These observations point out differences in early pathological events despite similar IOP elevation. In *Tg.CreMYOC^Y^*^437^*^H^*mice, elevated IOP may cause primary damage to ONH resulting in loss of transport, which then proceeds to loss of RGCs. In MB-model, elevated IOP may cause direct loss of RGC soma, which then progresses to axonal degeneration and impaired axonal transport. Several lines of evidence support that ONH is the first site of injury due to mutant MYOC-induced OHT. First, significant loss of RGC axons was observed as early as 10 weeks while RGC soma loss was observed later at 15 weeks. Second, VEP, which measure post-retinal function of visual system (including ON and visual centers of the brain) is reduced significantly prior to RGC functional or structural loss. Third, impaired axonal transport was observed in the ONH region as early as 7-weeks of injection prior to RGC functional loss in age-independent manner.

Both microtubules and neurofilaments are key cytoskeleton proteins required for axonal transport and loss of these proteins is associated with glaucomatous neurodegeneration^63, 65, 66^. Here, we propose that loss of microtubule and neurofilament at the ONH due to chronic IOP elevation is likely to responsible for impaired axonal transport. Both TEM and immunostaining demonstrated dramatic loss of cytoskeleton proteins including microtubules and neurofilament associated with impaired axonal transport. This is the first study showing that IOP elevation induces loss of microtubules impairing axonal transport in the ONH prior to neuronal loss. Since both microtubules and neurofilament assembly is regulated by autophagy ^67^, it is possible that chronic IOP elevation leads to compromised autophagy, which results into loss of neurofilament and microtubule assembly accumulating defective organelles in the ONH region. Consistent with this, several studies have suggested that chronic IOP elevation leads to compromised autophagy in the RGC axons^68, 69^. Alternatively, loss of microtubules, which facilitate formation and transport of autophagic vesicles carrying defective organelles may contribute to accumulation of autophagic vesicles in the ONH region including defective mitochondria as evident in our studies. Further understanding of these early events of axonal degeneration provides a therapeutic window for regeneration strategies to support dying axons.

In summary, we developed a Cre-inducible mouse model of POAG that faithfully replicates all features of human POAG including IOP elevation due to TM dysfunction and IOP-dependent glaucomatous neurodegeneration. We expect that *Tg.CreMYOC^Y^*^437^*^H^* mouse model will be a valuable tool to study the pathophysiology of glaucomatous damage to TM and RGC axons and provide new targets for the treatment.

## Methods

### Animal husbandry

Animals were housed and bred in a standard 12-hour light/12-hour dark conditions. They were fed standard chow *ad libitum* and housed in cages with dry bedding. Two-to three-month-old C57BL/6J mice (including both males and females) were obtained from the Jackson Laboratory (Bar Harbor, ME, USA). mT/mG mice were obtained from Jackson Lab (stock # 007576) and bred as described previously^38^. *Tg.CreMYOC^Y^*^437^*^H^* transgenic mice on pure C57BL/6J background were generated in the lab as described below. Mice were euthanized via inhalation of carbon dioxide followed by cervical dislocation as described previously ^33^.

### Generation of *Tg.CreMYOC^Y^*^437^*^H^* mice

*Tg.CreMYOC^Y^*^437^*^H^*mice were generated using the TARGATT site-specific transgenic technology (Applied Stem Cell) according to the manufacturer’s protocol^31^. Briefly, the cDNA for MYOC with the Y437H mutation was subcloned into a TARGATT plasmid (AST-3050, Applied Stem Cell) that contains the attB recombination site. The expression cassette contained a STOP signal flanked with a loxP site in the front of the Y437H MYOC cDNA under the control of CAG promoter. A mixture of the donor plasmid (containing the attB sites and the transgene) and the ΦC31 mRNA was injected into the pronuclei of H11P3-C57BL/6 mouse embryos. Genomic DNA of a founder was analyzed using primer pairs for the transgene and H11 locus to verify site-specific insertion. The primer set; AST 2005 (Applied Stem Cell, Inc) was utilized to detect H11 site specific integration. Transgene positive mice were bred with C57BL/6J mice and heterozygous mice were utilized for experiments. The following primers were utilized to detect the transgene during regular genotyping. Forward primer: CTCAGCAGATGCTACCGTCA, and reverse primer: GCACCTTGAAGCGCATGAA.

### Cre recombinase induction by viral vectors

Ad5-Cre or HAd5-Cre under the control of CMV promoter was utilized to selectively induce Cre activity in the TM. Ad5 and HAd5 expressing empty cassettes were utilized as controls. These vectors were purchased from the Viral Core Facility at the University of Iowa. Mice were anaesthetized using 2.5% isoflurane plus 100% oxygen. A single intravitreal injection of Ad5 or HAd5 expressing Cre, or empty cassette (2 × 10^7^ pfu/eye) was performed as described previously^32, 33, 70, 71^. Viral injections were performed in 3 to 6 months old male and female *Tg.CreMYOC^Y^*^437^*^H^*mice in all experiments except for axonal transportation in which 15-month-old *Tg.CreMYOC^Y^*^437^*^H^*mice were utilized. For CTB injections in 15-month-old *Tg.CreMYOC^Y^*^437^*^H^* mice, we have utilized intracameral injection of HAd5-empty or Cre as described previously^72^. For all other experiments, intravitreal injection was utilized.

### Antibodies and reagents

The following antibodies were used in the current study. Myocilin monoclonal antibody (catalog # 60357-1-Ig, Proteintech), DsRed antibody (catalog # 600-401-379, Rockland), KDEL (catalog # Ab12223, Abcam), ATF4 (catalog # SC-200, Santa Cruz Biotechnology)^36^, CHOP (catalog # 13172, Novus Biologicals), GRP78 (catalog # ab21685, Abcam), GRP94 (catalog # 11402, Santa Cruz Biotechnology), RBPMS (catalog # 118619, Gene Tex), GAPDH (catalog # 3683, Cell Signaling Technology), TOM20 (catalog # 11802-1-Ap, Proteintech), TUJ1(catalog # GTX130245, Genetex), GFAP (catalog #, ab4674-1001, Abcam), Fibronectin (catalog # AB2413-1001), Neurofilament H (catalog # 80170l, Biolegend), Phalloidin stain for actin (catalog #12956, Cell Signaling), and anti-alpha SMA (catalog # ab5694, Abcam).

### Real-time PCR

The anterior segment of Cre^-^ and Cre^+^ *Tg.CreMYOC^Y^*^437^*^H^* mice was submerged in RNAprotect Cell Reagent (catalog #76526, QIAGEN). Subsequently, the anterior segment was pelleted by centrifugation for 5 minutes at 600g and then subjected to total RNA extraction using a RNeasy Mini Kit (QIAGEN). Following RNA extraction, real-time PCR was carried out using the SYBR® Green Quantitative RT-qPCR Kit (catalog # QR0100, Sigma Aldrich) with the following primers: Forward primer, 5’-TGACTTGGCTGTGGATGAAG-3’; Reverse primer: 5’-TTGTCTCCCAGGTTTGTTCG-3’.

### Western blot analysis

Ocular tissues were carefully dissected and lysed in RIPA buffer (Thermo Fisher) as described previously^30, 36^. Approximately 20–30 μg of total protein was separated on denaturing 4–12% gradient polyacrylamide ready-made gels (NuPAGE Bis–Tris gels, Life Technologies, Grand Island, NY, USA) and subsequently transferred onto PVDF membranes. The blots were blocked with 10% non-fat dried milk for 1 hour and then incubated overnight with specific primary antibody at 4 °C on a rotating shaker. Membranes were washed three times with phosphate-buffered saline (PBS)/Tween buffer (PBST) and were incubated with the corresponding HRP-conjugated secondary antibody for 90 minutes. Proteins were visualized on the LI-COR Odyssey Fc image system using ECL detection reagents (Super Signal West Femto Maximum Sensitivity Substrate; Life Technologies)^32, 47^. The same blot was subsequently incubated with a GAPDH antibody (Cell Signaling Technology Inc) to ensure the equal protein loading.

### Anterior segment flat mount

Whole anterior segment flat mount was performed to image DsRed-MYOC levels in the entire TM region. Cre^-^ and Cre^+^ *Tg.CreMYOC^Y^*^437^*^H^* mice were enucleated, fixed with 4% paraformaldehyde (PFA), and dissected carefully to separate the anterior and posterior segment. Anterior segments were then transferred onto a slide and cut into four quadrants to place tissue flat on the slide. Anterior segments were mounted with DAPI-mounting solution and images were captured using a Keyence microscope (Itasca, IL, USA).

### RNA scope

Fluorescent *in* situ hybridization was performed using the RNAscope^®^ Multiplex Fluorescent Assay v2 (ACD Diagnostics) following modifications as described before ^73^. Briefly, fresh frozen histologic sections of mouse eyes were pretreated according to the manufacture’s protocol using hydrogen peroxide and target retrieval reagents including protease IV. Probes were then hybridized according to the protocol and then detected with TSA Plus^®^ Fluorophores fluorescein, cyanine 3, and cyanine 5. Sections were mounted with Prolong Gold Antifade (Thermo Fisher) with coverslip for imaging and imaged using confocal microscope (SP8, Leica). Probes specific for *Myoc* (Cat # 506401-C3) and *MYOC* (Cat # 506541) transcripts were designed by the manufacturer.

### Immunostaining

Enucleated eyes were fixed with 4% PFA for 3 hours. The eyes were embedded in paraffin or OCT compound (Tissue-Tek). Sections of 5 or 10 microns were cut and utilized for immunostaining as described previously^30, 47^. For paraffin-embedded samples, the slides were deparaffinized and rehydrated, followed by antigen retrieval with citrate buffer (pH = 6). For OCT-embedded sections, slides were washed with PBS. The slides were then incubated with blocking buffer (10% goat serum and 0.5% Triton-X-100 in PBS) for 2 hours. The slides were incubated with the primary antibody in blocking buffer overnight. After three washes with PBS, the slides were incubated with the appropriate Alexa Fluor secondary antibodies (Life Technologies, Grand Island, NY, USA) for 2 hours. Following three final washes in PBS, the sections were mounted with DAPI-mounting solution, and images were captured using a Keyence microscope (Itasca, IL, USA) as described previously ^42, 47^.

### Mouse slit-lamp examination

To evaluate ocular abnormalities and inflammation in HAd5-Empty and HAd5-Cre-injected eyes, slit-lamp microscopy (SL-D7; Topcon) was performed as described previously^30^.

### IOP measurements

IOP measurements were conducted using the TonoLab® rebound tonometer (Colonial Medical Supply, Franconia, NH, USA) under anesthetic conditions (isoflurane (2.5%); oxygen (0.8 L/min)) as described previously ^36^ All IOP assessments were performed between 10 AM and 2 PM in a masked manner. In addition, IOP measurements were replicated by another technician in a masked manner. The final IOP value was obtained by averaging six individual IOP measurements. Conscious IOPs were measured as described previously^74^.

### Aqueous humor outflow facility

The outflow facility in Cre^-^ and Cre^+^ *Tg.CreMYOC^Y^*^437^*^H^* mice was assessed using the constant-flow infusion method as previously described^74^ Briefly, mice were anesthetized with a ketamine/xylazine solution (100/10 mg/kg), and body temperature was maintained at a physiological level using a 37°C electric heating pad throughout the procedure. Topical ocular proparacaine HCl (0.5%) was administered to induce corneal anesthesia. Anterior chambers were cannulated with 32-gauge ½” steel needles (Steriject, Keeler, USA) through the cornea without contacting the iris, anterior lens capsule, or corneal endothelium. The needles were connected to tubing running to a pressure transducer. The opposite end of the transducer was connected by more tubing to a 100 μL microsyringe (Hamilton Co., Reno, NV, USA) loaded into a microdialysis infusion pump, which was filled with sterile PBS. Eyes were infused at a flow rate ranging from 0.1 μL/min to 0.5 μL/min (in 0.1 μL/min increments), and three stabilized pressures at 15-minute intervals were recorded for each flow rate. The aqueous humor outflow facility was then calculated as the reciprocal of the slope (determined by simple linear regression) of the linear part of a plot of mean stabilized pressure as the ordinate and flow rate as the abscissa. In a small proportion of cases (13%), data points obtained at flow rates of 0.5 μL/min deviated from linearity. Such points were excluded from the analysis.

### Pattern electroretinography (PERG)

The RGC function was assessed using binocular snout pattern electroretinography (PERG) system (JORVEC Corp., Miami, FL, USA) as described previously^47^. Briefly, mice were anesthetized with an intraperitoneal injection of ketamine/xylazine mixture (100, 10 mg/kg, respectively). Anesthetized mice were positioned on a temperature-controllable metal base at a fixed distance (10 cm) from LED monitors and maintained at a constant body temperature (37°C) using a rectal probe. To prevent corneal dryness during recording, a small amount of hypromellose eye drops was applied topically^33, 47^. The PERG was simultaneously derived from each eye using subcutaneous electrodes placed at the snout (active), back of the head (reference), and tail (ground) in response to contrast reversal of gratings generated from two LED screens operating at slightly different frequencies. Two consecutive readings were averaged, and amplitudes (P1–N2) and latencies indicating RGC soma function were shown graphically.

### Whole-mount retina staining with RBPMS

Total number of RGCs was analyzed using whole-mount retina staining with the RBPMS antibody as described previously^47^. The enucleated eyes were fixed with 4% PFA for 12 hours at 4 °C. After rinsing the eyes with PBS, the anterior segment was removed, and the retinas were carefully separated from the posterior cup. The isolated retinas were incubated with blocking buffer (PBS containing 10% goat serum and 0.2% Triton X-100) for 12 hours at 4 °C. Subsequently, the retinas were incubated with the RBPMS antibody for 3 days at 4 °C followed by a 2-hour wash in PBS. After washing, the retinas were incubated with a corresponding secondary antibody (goat anti-rabbit 568, 1:500; Invitrogen) for 2 hours at room temperature, washed with PBS three times, and then mounted with a mounting medium containing DAPI nuclear stain. For RGC counting, a minimum of 16 non-overlapping images from the entire retina were captured at 200x magnification using a Keyence fluorescence microscope (Itasca, IL, USA), and RBPMS-positive cells were counted using ImageJ software^32, 33, 47^.

### Assessment of optic nerve degeneration

Optic nerve axonal degeneration was assessed using a paraphenylenediamine (PPD) staining as described previously^33, 47^ In brief, optic nerves were fixed overnight in PBS containing 3% glutaraldehyde/paraformaldehyde at 4°C. The optic nerves were rinsed twice for 10 minutes with 0.1M phosphate buffer and once with 0.1M sodium acetate buffer. Optic nerves were then dehydrated in graded ethanol concentrations. After embedding in resin, transverse semithin sections (1 μm) were cut and stained with 1% PPD for 10 minutes. Ten images without overlap were captured using confocal Leica microscope and counted manually using Image J software. The sum of total surviving axons counted approximately equaled to 10% of the total optic nerve cross-sectional area.

### Transmission electron microscopy (TEM)

To perform TEM imaging of the outflow pathway, 1 μm sections were processed for TEM analysis as described previously^47^ The animals were perfusion-fixed with 4% PFA, and eyes were enucleated. The anterior chamber and optic nerve were dissected carefully and further fixed with 1.5% paraformaldehyde and 1.5% glutaraldehyde in 100 mM sodium cacodylate buffer (pH 7.2) for at least 24 hours. After aldehyde fixation, the eyes were incubated with 1% osmium tetroxide in 100 mM sodium cacodylate buffer, followed by ethanol dehydration and embedding of the samples in Eponate 812 resin. Sections (1 μm) were cut using an ultramicrotome (EM UC6; Leica) and stained with uranyl acetate and Reynolds lead citrate. Images were taken with a transmission electron microscope (JEM-1230; JEOL) equipped with a 2K × 2K CCD camera (USC1000; Gatan Inc.) at the University of Iowa Central Microscopy Research Facility.

### Visual electrode potential

Mice were dark adapted for eight hours in a light-proof box. The dark-adapted animals were anaesthetized with an intraperitoneal injection of a ketamine/xylazine mixture (100, 10 mg/kg, respectively). Pupils were dilated using a single drop of 0.5% tropicamide for five minutes. Visual evoked potentials (VEPs) were recorded using the Celeris D430 rodent ERG system (Diagnosys LLC, MA, USA) under maintained dark conditions as described previously^75^. Mice were positioned on a heated platform to regulate body temperature, and corneal moisture was maintained with 1-2% hydroxypropyl methylcellulose. Corneal electrodes integrated with stimulators were applied to lubricated corneas. Ground and reference needle electrodes were subcutaneously inserted into the tail, cheek, and subdermal region of the head, approximating the visual cortex. The midpoint between the ears served as the reference point. Although recording locations were not histologically verified, they were assumed to be consistent based on an impedance of 1.5-10 kΩ. Each eye was subjected to 100 flashes of 1 Hz, 0.05 cd s/m² white light delivered through corneal stimulators. Recordings were captured for 300 ms at a sampling rate of 2000 Hz. Two trials consisting of 50 sweeps were performed for each mouse. VEP data was filtered with low and high frequency cutoffs of 1.25 Hz and 100 Hz, respectively. The amplitude of N1 was calculated as the distance from baseline to the most pronounced negative peak, while P1 amplitude was determined from the deepest negative peak to the subsequent positive peak. Latency was defined as the interval between stimulus onset and the initial response peak^76^.

### Axonal transport

To assess axonal anterograde transport mechanism, fluorescently labeled cholera toxin B (CTB) was injected intravitreally and its transport across ON to the SC was tracked using fluorescence microscope as described previously^47^. Both 4- and 15-month-old *Tg.CreMYOC^Y^*^437^*^H^* mice were used to determine whether CTB transport blockage is age-dependent, as described by a previous study^49^. Seven weeks after either HAd5-empty or HAd5-Cre injection, *Tg.CreMYOC^Y^*^437^*^H^* mice were anesthetized using isoflurane and an intravitreal injection of 3 µL of 0.1% CTB (reconstituted in PBS) conjugated with either Alexa Fluor 555 or Alexa Fluor 480 (Invitrogen, Life Technologies, Grand Island, NY, USA) was performed. Forty-eight-hours later, mice were euthanized, and eyes and whole brains were fixed in 4% PFA at 4°C for 12 hours. Subsequently, tissues were washed with PBS, cryoprotected in a sucrose gradient (10-30%), embedded in OCT compound, and cryo-sectioned at a thickness of 10 µm. Axonal anterograde transport of CTB to ON and SC was directly visualized using fluorescence microscopy. Tissue sections were washed with PBS, mounted with a DAPI-containing mounting medium, and imaged using a Keyence fluorescence microscope^47^.

### Microbead occlusion model

Microbead occlusion model was developed by injecting magnetic beads in the anterior chamber. The magnetic beads block TM outflow, elevating IOP and inducing RGC loss as described previously^21, 77^. The experiment was performed in 4-month-old male C57BL/6J mice. Anesthetized animals were given topical eye drops of Tropicamide (0.5%, one drop) to dilate the pupil and a drop of local anesthetic (0.5% proparacaine hydrochloride) was applied to the cornea. Angled forceps were used to proptose the eye. The cornea was then breached with a glass pulled micropipette that is guided to the anterior chamber. One to 1.5 microliters of 5.8 µm diameter magnetic microbeads (Bangs Laboratories, Fishers, IN) were injected into the anterior chamber of both eyes using a manual micromanipulator (WPI, Sarasota, FL). A magnet was used to draw the beads in place before withdrawing the micropipette. Injected eyes were then covered in ophthalmic ointment and mice were placed on a heating pad for recovery. Another group of mice that received PBS served as controls. IOP was measured every week to monitor IOP elevation. We observed stable IOP elevation of more than 4mmHg or more in 50-60% of eyes injected with microbeads likely due to failure of the procedure to keep magnetic beads throughout the TM. We therefore selected mice that develop OHT and utilized them for subsequent PERG analysis, RBPMS staining and axonal transport analysis.

### Study approval

Animal studies were performed in agreement with the guidelines of the Association for Research in Vision and Ophthalmology (ARVO) Statement for the Use of Animals in Ophthalmic and Vision Research. Experimental protocols were approved by appropriate IACUC and Biosafety Committees at the University of California, Irvine (UCI).

### Statistics

Prism 9.0 software (GraphPad, San Diego, CA, USA) was used for statistical analyses. Data was represented as mean ± SEM. For all experiments, n refers to the number of eyes. *P < 0.05* was considered statistically significant. The student’s *t*-test was used to compare 2 groups. For comparison of different treatments, 2-way ANOVA was used followed by a Bonferroni post-hoc correction.

## Authors Contributions

GSZ, and BK designed research studies. BK, RK, and LL, performed experiments and analyzed data. JCM, SY, YS, PM, KP, WC and DSK assisted in some experiments. BK and GSZ wrote the manuscript. KP assisted in editing of the manuscript. All authors discussed the results and implications and commented on the manuscript at all stages.

## Acknowledgments

These studies were supported by the National Institutes of Health (EY034333 and EY026177). The authors acknowledge support from NIH grant EY034238 and from an unrestricted grant from Research to Prevent Blindness to the Gavin Herbert Eye Institute at the University of California, Irvine.

## Data Availability Statement

All the datasets used and/or analyzed in the present study are available from the corresponding author on reasonable request.

