## Supplementary material for "Impaired axonal transport at the optic nerve head contributes to neurodegeneration in a novel Cre-inducible mouse model of myocilin glaucoma": SI

**Supplemental Information**

**Kaipa *et al*.**

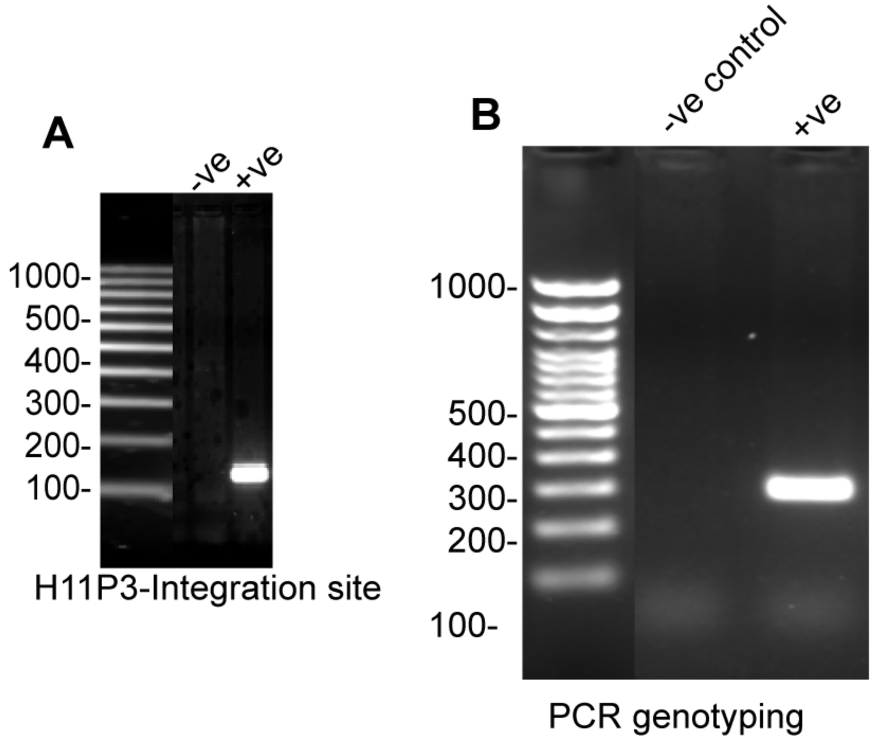


**Figure SI-1. Confirmation of transgene integration at the H11 site in *Tg.CreMYOC*$^{Y437H}$ mice**. (**A**) To confirm a site-specific knock-in of the transgene at the H11 site, PCR was performed, using primers specific to the integration site. PCR demonstrated a stable integration of transgene in one founder line. This founder was further bred with C57BL/6J mice, and offspring were utilized for subsequent studies. (**B**) To perform the regular genotyping for the presence of the transgene, PCR primers were utilized, which recognize human MYOC and the DsRed Region. The positive PCR band at 300 base pairs indicated the presence of the transgene.

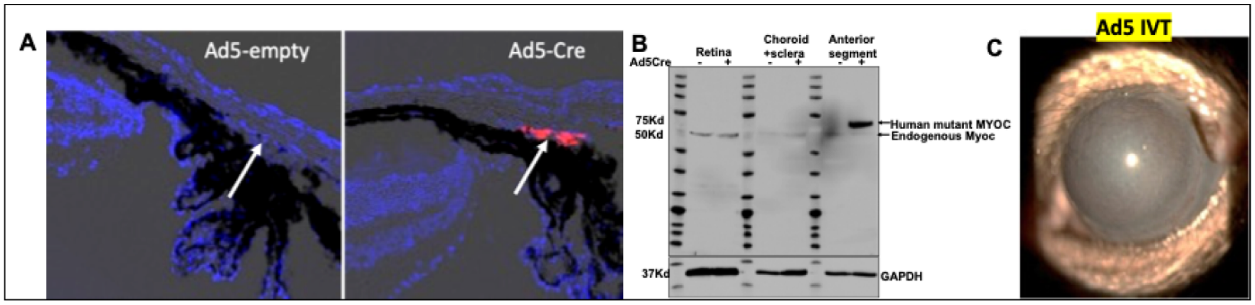


**Figure SI-2. Ad5-Cre induces mutant myocilin in the TM cells of *Tg.CreMYOC$^{Y437H}$* mice.** *Tg.CreMYOC$^{Y437H}$* mice were injected intravitreally with Ad5-empty or Ad5-Cre (2x10*7pfu/eye). (**A**) DsRed (indicating mutant myocilin) was localized to the TM cells of *Tg.CreMYOC$^{Y437H}$* mice, 5-weeks after injection of Ad5-Cre (n=6 mice); no DsRed expression was found in other ocular tissues. (**B**) Various ocular tissues were harvested from *Tg.CreMYOC$^{Y437H}$* mice, 5-weeks after injection of Ad5-empty (control) or Ad5-Cre; aliquots of the tissue samples were subjected to Western-blot analysis for myocilin. The Western blots showed the presence of 75-kDa human mutant myocilin exclusively in iridocorneal angle tissues from the Ad5-Cre-injected eyes, whereas endogenous WT myocilin was detected in all of the ocular tissues, using the myocilin antibody. (**C**) Slit lamp image of mice, 5-weeks after injection with Ad5-Cre, showing that ocular inflammation is associated with Ad5-Cre.

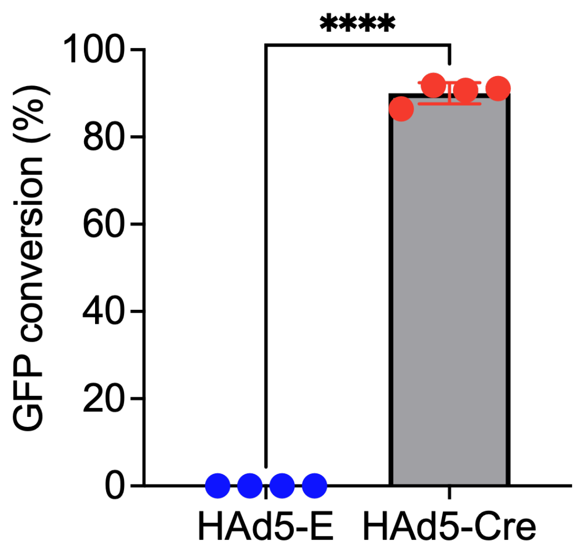


**Figure SI-3. HAd5-Cre transduces robust Cre activity in mT/mG fluorescence-based reporter mice.** mT/mG reporter mice were injected intravitreally with HAd5-Cre (2 × 10^7 pfu/eye), and the conversion from tdTomato to GFP was examined after one week, using confocal microscopy (n=4). GFP-positive TM cells were counted, and their number was divided by the total number of TM cells present in each section (based on DAPI); the fraction X 100 is represented as % GFP conversion. Over 90% of the TM cells were transduced by HAd5-Cre injection compared to HAd5-E injection ($P < 0.0001$, unpaired two-tailed t-test; mean ± SEM).

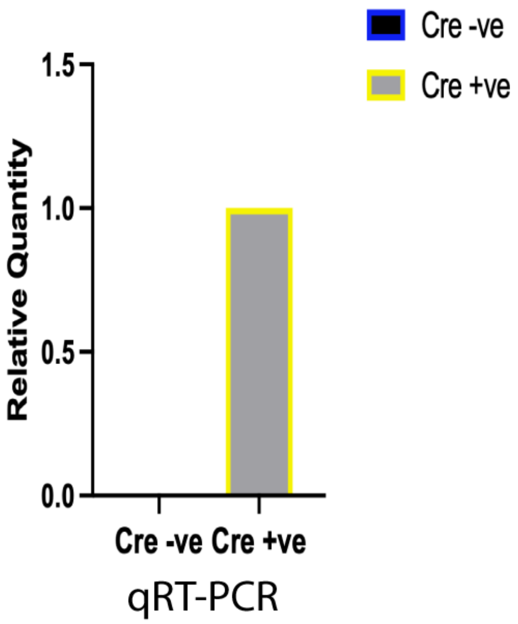


**Figure SI-4. An intravitreal injection of HAd5-Cre recombinase induces mutant MYOC transcript in the anterior segment of *Tg.CreMYOC$^{Y437H}$* mice.** HAd5-empty or HAd5-Cre were injected intravitreally (2x10^7 pfu/eye) into *Tg.CreMYOC$^{Y437H}$* mice, and samples of the isolated anterior segments from the respective mice were subjected to QPCR, using primers specific to human MYOC. Quantitative analysis of *MYOC* mRNA expression levels using qPCR indicated induction of MYOC mRNA exclusively in HAd5-Cre-injected mice ("Cre +ve"). MYOC transcript was normalized to GAPDH.

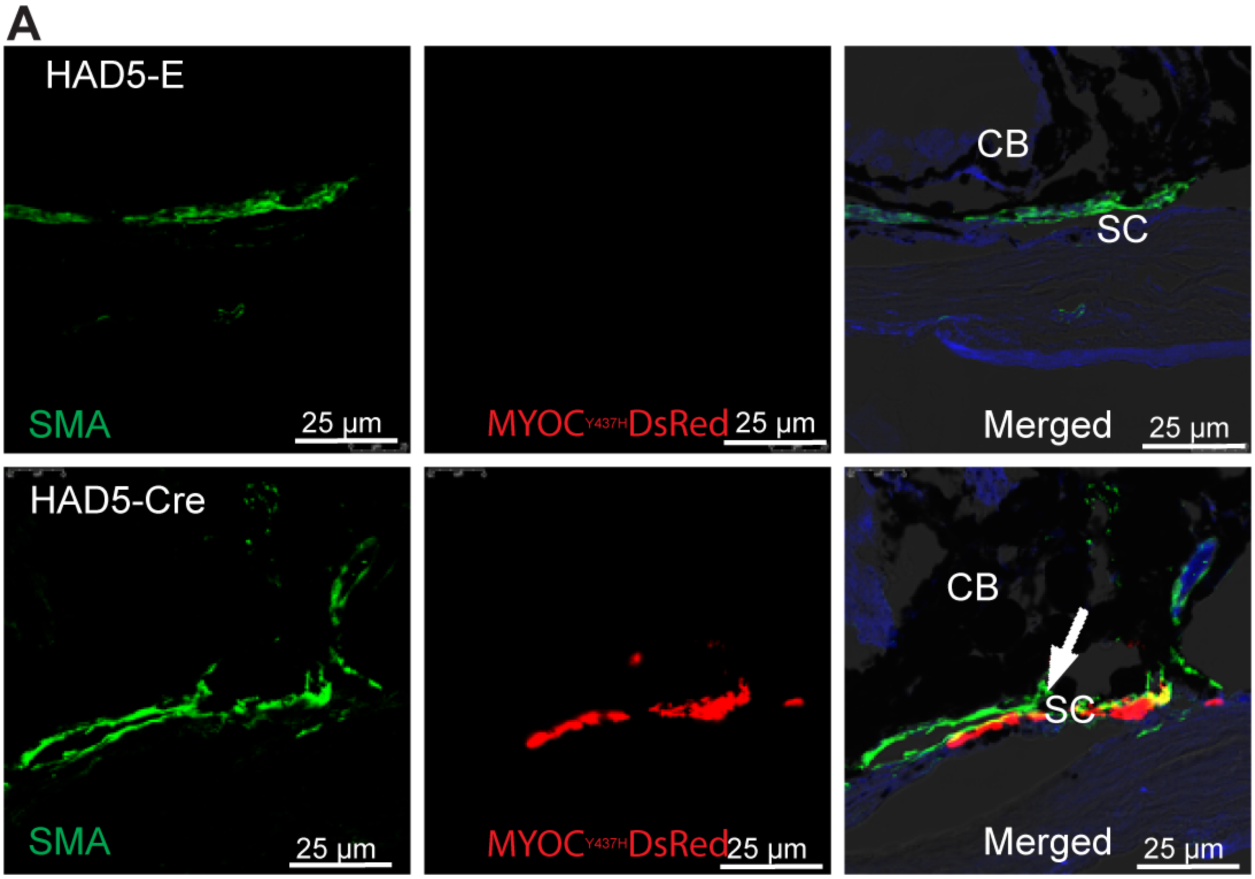


**Figure SI-5. Mutant myocilin colocalizes with the TM-specific marker, smooth muscle actin (SMA) in Cre$^{+}$*Tg.CreMYOC$^{Y437H}$*** mice**.** Anterior-segment cross sections from Tg.Cre.MYOC$^{Y437H}$ mice, injected with HAd5-empty or HAd5-Cre, were immunostained with antibody against alpha smooth muscle actin (SMA), which predominantly stains TM and ciliary muscle. We observed strong co-localization of SMA with DsRed-tagged mutant MYOC in the TM region but not in the ciliary muscle region, indicating that mutant myocilin is specifically expressed in the TM.

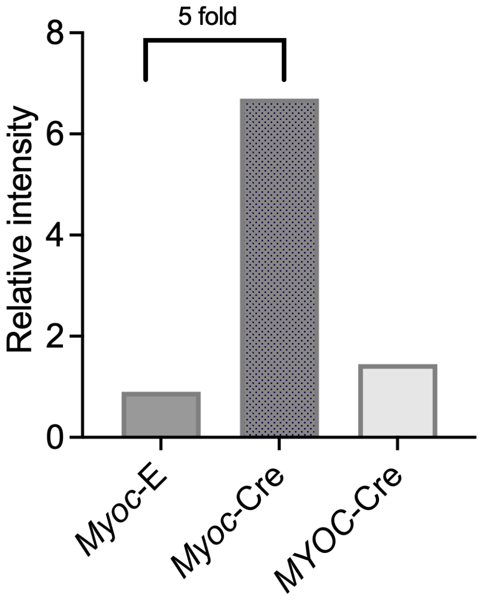


**Figure SI-6. Mutant *MYOC* is expressed at a level comparable to endogenous *Myoc* in *Cre⁺Tg.Cre.MYOC*$^{Y437H}$ mice.** mRNA transcripts of mutant *MYOC* and endogenous *Myoc* from *Tg.Cre.MYOC*$^{Y437H}$ mice, injected with HAd5-empty or HAd5-Cre, were analyzed through RNA scope, and images were captured using confocal microscopy. Integrated fluorescence intensities of mutant *MYOC* and endogenous *Myoc* were analyzed using Image J. software. We observed that mutant *MYOC* is expressed at a level similar to endogenous *Myoc*. Expression of mutant MYOC induced endogenous *Myoc* expression by 5-fold in *Tg.Cre.MYOC*$^{Y437H}$ mice injected with HAd5-Cre. Myoc-E represents endogenous Myoc transcripts from HAd5-empty-injected *Tg.Cre.MYOC*$^{Y437H}$ mice. Myoc-Cre and MYOC-Cre represent endogenous Myoc and mutant MYOC transcripts, respectively, from HAd5-Cre-injected *Tg.Cre.MYOC*$^{Y437H}$ mice.

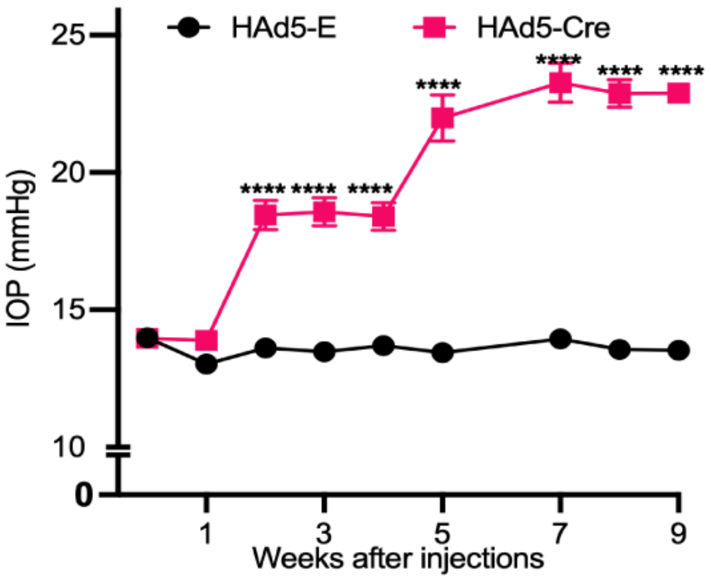


**Figure SI-7. Conscious IOP measurements in Cre$^{-}$ and Cre$^{+}$ *Tg.CreMYOC$^{Y437H}$* mice**
Conscious IOP (without the use of anesthesia) was measured, in a masked manner, in *Tg.CreMYOC$^{Y437H}$* mice each week from 1 to 9 weeks after intravitreal injection of Had5-empty (control) or Had5-Cre. A significant IOP elevation was observed in HAd5-Cre-injected mice compared to control, starting at 2-weeks after injection. Data are presented as mean ± SEM (*n* = 8 for HAd5-E; *n* = 10 for HAd5-Cre), analyzed by 2-WAY ANOVA with multiple comparisons, (*****P* < *0*.0001).

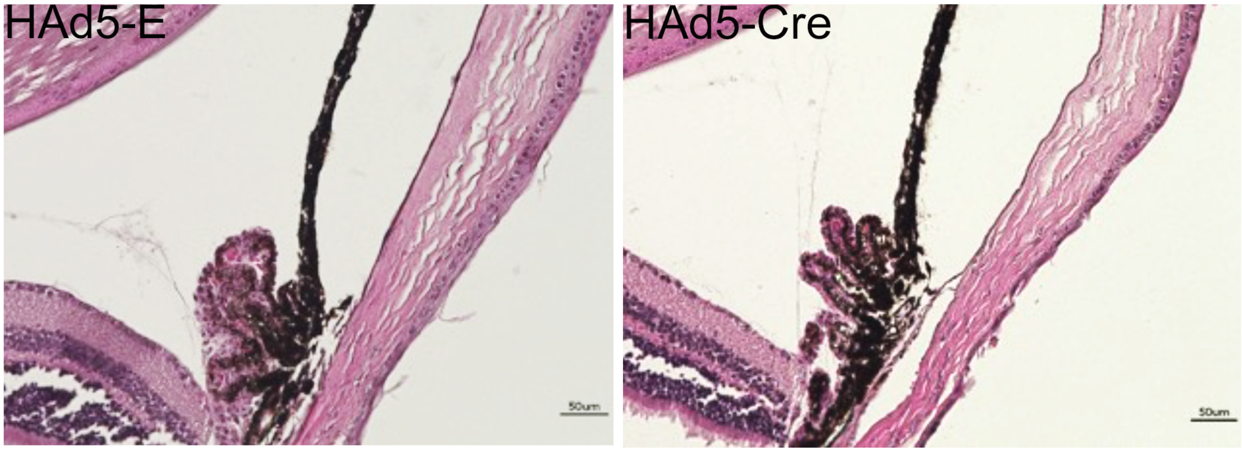


**Figure SI-8. Open iridocorneal angle, and no gross abnormalities in *Tg.CreMYOC$^{Y437H}$* mice.** *Tg.CreMYOC$^{Y437H}$* mice were injected with HAd5-empty or Had5-Cre, and *H and E* staining was performed at 5-weeks post-injection. Histological analysis demonstrated open angle and normal ocular structures in both groups of mice (n=3).

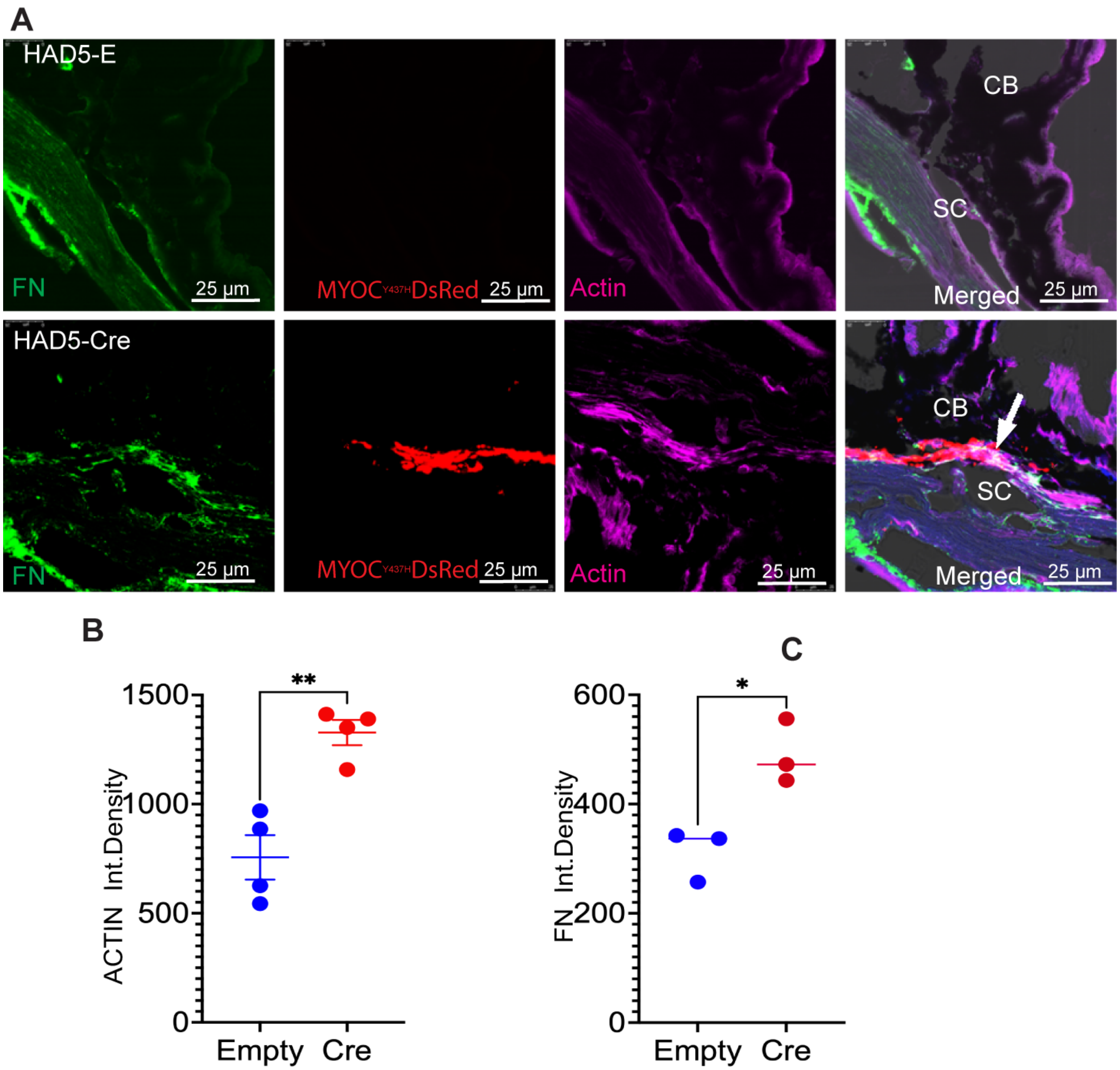


**Figure SI-9. Mutant myocilin expression is linked to increased fibronectin and actin in the TM *Cre$^+$Tg.CreMYOC$^{Y437H}$* mice.** Anterior segments from Tg.CreMYOC$^{Y437H}$ mice injected with Had5-empty (Cre$^-$, control) or Had5-Cre (Cre$^+$) were immunostained for MYOC (Ds-Red), actin (phalloidin; purple), and fibronectin (green). (**A**) Representative images for control mice (upper panels) and Cre$^+$-mice (lower panels) demonstrated increased fibronectin and actin labeling, which strongly co-localized in the TM (right-most panels). Quantification of integrated intensity for actin (**B**) and fibronectin (**C**) demonstrated a significant increase in actin (**P = 0.0028, unpaired two-tailed t-test;

mean ± SEM, n = 4) and fibronectin (*P = 0.0149, unpaired two-tailed t-test; mean ± SEM, n = 3) levels in Cre$^+$Tg.CreMYOC$^{Y437H}$ mice relative to the controls.

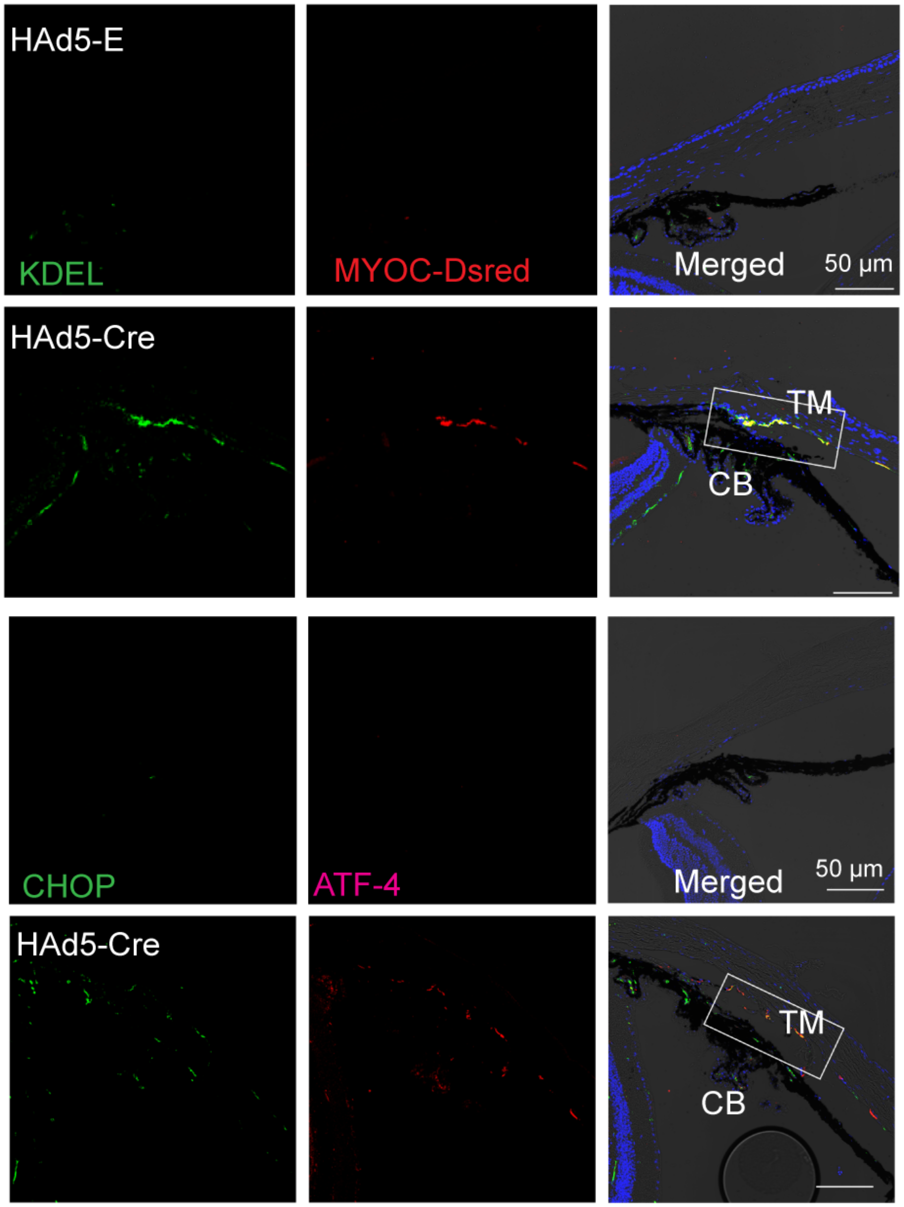


**Figure SI-10. Mutant myocilin expression in mouse TM induces ER stress**. Anterior segments from *Tg.CreMYOC*$^{Y437H}$ mice, 5-weeks after injection with HAd5-empty or Had5-Cre were immunostained for various ER stress markers, including KDEL (Top panel) and CHOP+ATF4 (Bottom panel). KDEL (which recognizes GRP78 and GRP94), CHOP, and ATF4 were increased in the TM region of Cre-injected *Tg.CreMYOC*$^{Y437H}$ mice (n=3).

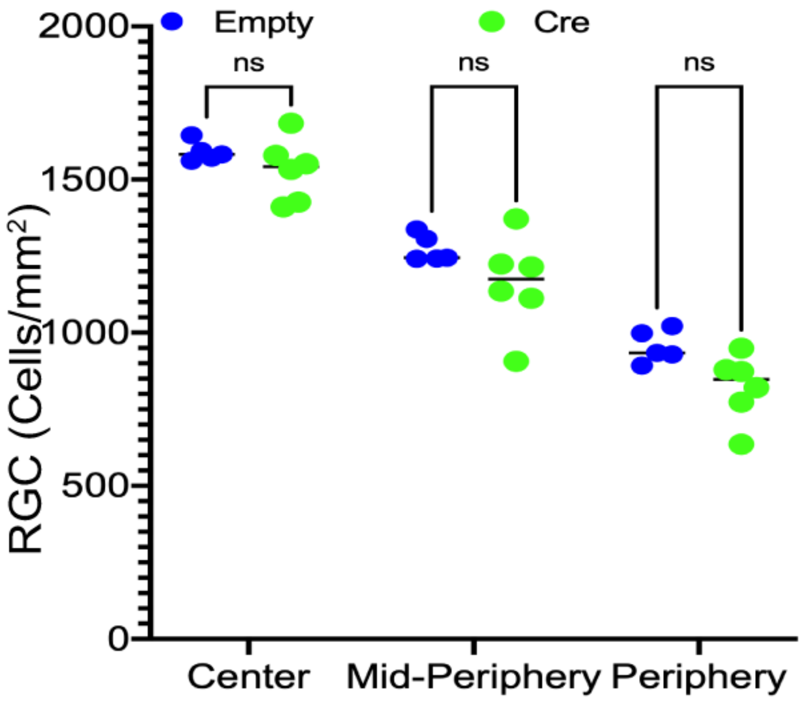


**Figure SI-11. RGC analysis in HAd5-Cre-injected *Tg.CreMYOC$^{Y437H}$* mice, 10-weeks after injection.** We examined whether HAd5-Cre injected *Tg.CreMYOC$^{Y437H}$* mice show RGC structural loss by performing RBPMS staining of whole mounts prepared from the mice at 10 weeks after injection. Analysis of individual RBPMS-positive RGCs did not show any significant RGC loss. (n=6, 2-WAY ANOVA with multiple comparisons).

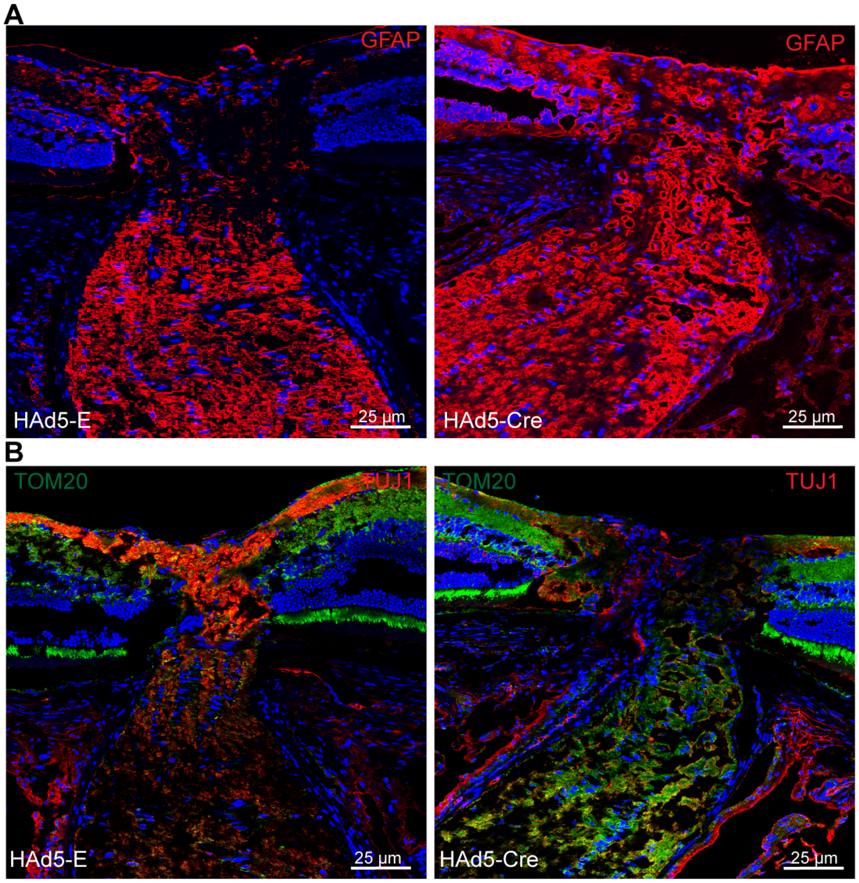


**Figure SI-12. IOP elevation leads to increased GFAP and loss of neuronal markers in Cre$^{+}$Tg.Cre.MYOC$^{Y437H}$ mice.** Retinal cross-sections were prepared from the eyes isolated from *Tg.CreMYOC$^{Y437H}$* mice 15-weeks after intravitreal injection with HAd5-empty or Had5-Cre, and immunostained to examine glial cell activation and/or loss of neuronal markers. **(A)** Immunostaining for GFAP demonstrated gliosis in the ONH region of Cre-injected *Tg.CreMYOC$^{Y437H}$* mice relative to controls (n=3 for both). (**B**) Immunostaining for TOM20 and Tuj1 (microtubule marker) demonstrated increased mitochondrial accumulation in ON axons of Cre-injected *Tg.CreMYOC$^{Y437H}$* mice relative to controls (n=3 for both).

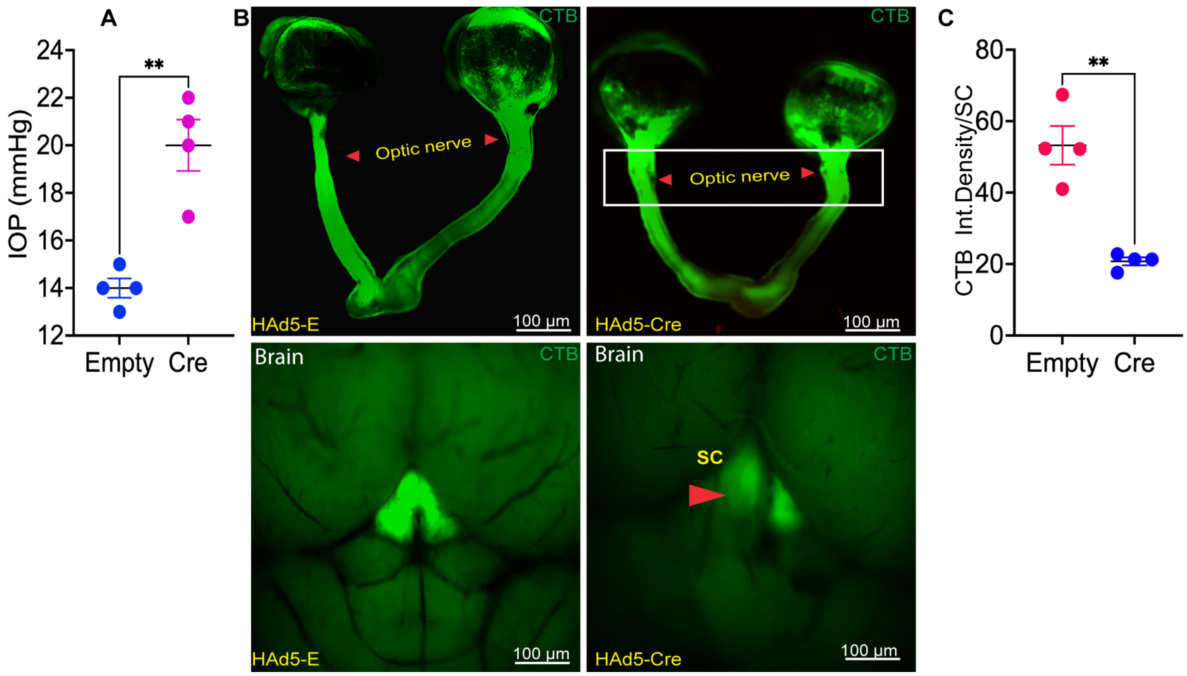


**Figure SI-13. Ocular hypertension induces anterograde transport deficits in young *Tg.CreMYOC$^{Y437H}$* mice.** Four-month-old *Tg.CreMYOC$^{Y437H}$* mice were intravitreally injected with either HAd5-Empty (control) or HAd5-Cre, and anterograde axonal transport mechanisms were assessed. (**A**) IOP measurements revealed significant IOP elevation in the HAd5-Cre mice, 6 weeks post-injection (n = 4; unpaired t-test; **P = 0.0020). At 7 weeks post-injection of HAd5-Empty or HAd5-Cre, the mice were then injected intravitreally with cholera toxin B (CTB) to determine the relative transportation of the CTB. (**B**) Representative images of CTB fluorescence are shown for the control mice (left panels) and Cre-injected mice (right panels). In the HAd5-Empty group (left), CTB was transported continuously along the entire length of the optic nerve to the superior colliculus (SC). In contrast, mice injected with Cre (right) displayed a significant transport block at the optic nerve head, and little CTB was detected in the SC. Fluorescence intensity of CTB in the SC was measured and represented graphically in (**C**), documenting 65% loss of CTB transportation to the SC of the Cre-injected mice (**P=0.00, n= 4 for HAd5-Empty, n = 3 for HAd5-Cre). (Note: CTB fluorescence analysis was performed with only those Cre-injected *Tg.CreMYOC$^{Y437H}$* mice in which the IOP was elevated by 4mmHg or more over the control.)

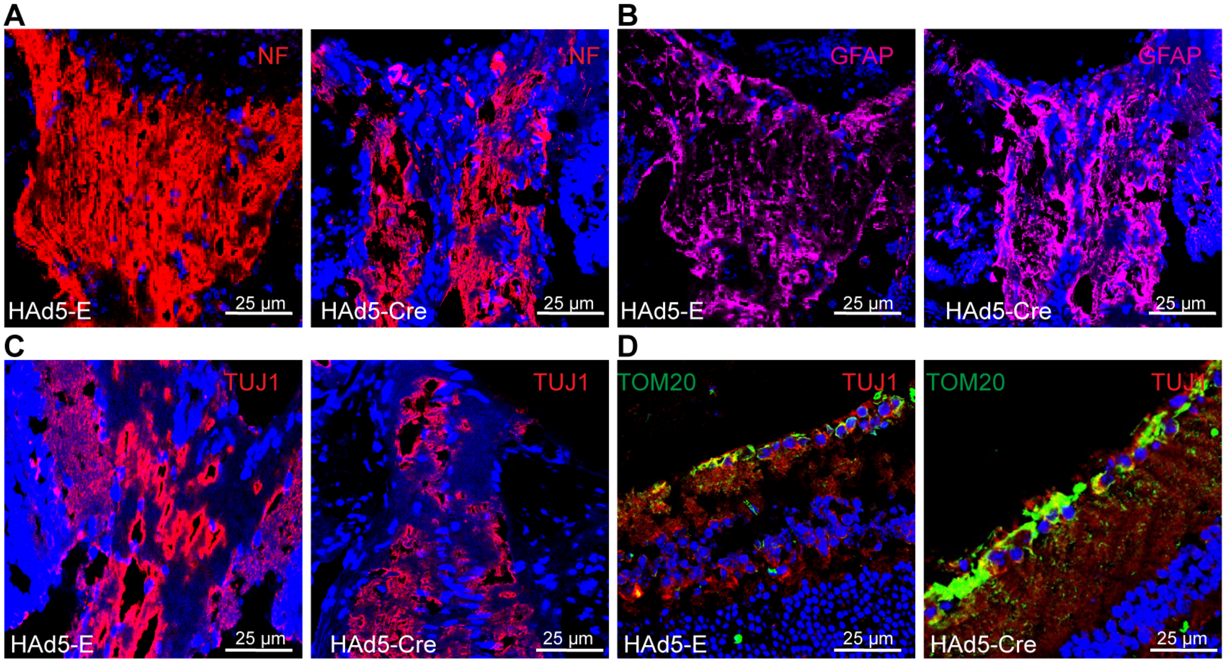


**Figure SI-14. Sustained IOP elevation leads to loss of microtubules and neurofilaments in the ONH of *Tg.CreMYOC$^{Y437H}$* mice.** To determine whether impaired axonal transport is associated with loss of cytoskeleton, we utilized *Tg.CreMYOC$^{Y437H}$* mice at 7 weeks post-injection of HAd5-Cre, which exhibited axonal-transport blockage. Immunostaining of retinal cross sections revealed reduced neurofilament levels (**A**) and diminished microtubule density (**C**), as indicated by TUJ1 antibody staining. Increased GFAP levels (B) indicated gliosis in the ONH region of the Cre-injected mice. These data indicate that loss of neurofilament and microtubules, which are required for active axonal transport is likely a cause for impaired axonal transport in the Cre$^{+}$ *Tg.CreMYOC$^{Y437H}$* mice. This impairment further results in abnormal accumulation of mitochondria in RGC soma as evident from increased TOM20 (**D**) at 12 weeks post-injection of HAd5-Cre (n=3).

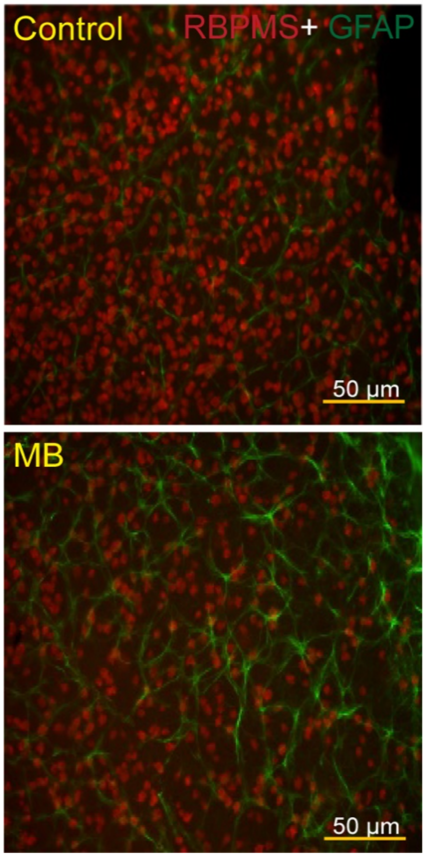


**Figure SI-15. Structural loss of RGCs in mouse model of microbead-injected mice.** Representative immunostaining for RBPMS (Red) and GFAP (green) in whole-mount retinas from mice, 6-weeks after injection of PBS or microbeads. Loss of RBPMS-positive RGCs along with increased GFAP reactivity was observed in the RGC layer of the whole mounts from the microbead-injected eyes.

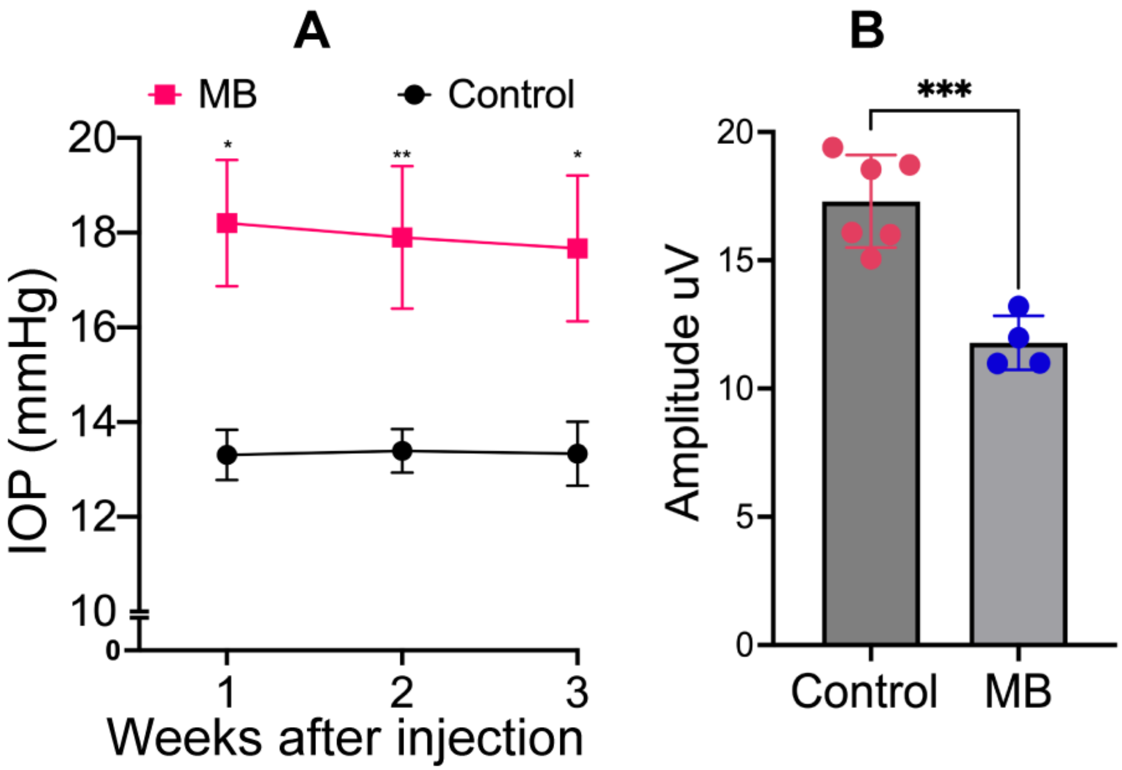


**Figure SI-16. Ocular hypertension and functional loss of RGCs in microbead-injected mice.** 4-month-old C57 mice were injected with PBS (control) or microbeads (MB) *via* the intracameral route, and IOP was monitored for 3 weeks. (**A**) IOP measurements revealed a significant IOP elevation in the MB-treated mice. Note that IOP is shown only for eyes in which the IOP was elevated by 4mmHg or more over the control (n=6 control, n=4 MB; 2-WAY ANOVA with multiple comparisons). (**B**) PERG analysis was performed on selected ocular hypertensive eyes to determine functional loss of RGCs. PERG analysis demonstrated a significant functional loss of RGCs (n=6 control, n=4 MB; unpaired t-test; ***P=0.0006). These mice were further subjected to analysis of CTB transportation. Note that these mice were further utilized for CTB transportation at 4-weeks post injection.
